# Lactuchelins: New lipopeptide siderophores from *Pseudomonas lactucae* inhibit *Xanthomonas campestris* pv. *campestris* 8004

**DOI:** 10.1101/2025.02.28.640635

**Authors:** Guillaume Chesneau, Alba Noel, Dimitri Bréard, Alice Boulanger, Martial Briand, Sophie Bonneau, Yujia Liu, Andrew Hendrickson, Torben Nielsen, Alain Sarniguet, David Guilet, Adam Arkin, Lauren Lui, Matthieu Barret

**Author notes:** G.C. and A.N. contributed equally to this work.

## Abstract

Seeds harbor diverse microbial communities, including beneficial microbes that play a vital role in protecting plants from seed-borne pathogens. Despite their critical importance, the molecular mechanisms driving intermicrobial competition within the seed microbiome remain poorly understood, limiting the potential to optimize seed inoculation strategies. In this study, we evaluated the inhibitory potential of 30 seed-borne bacterial strains against the phytopathogen *Xanthomonas campestris* pv. *campestris* 8004 (Xcc8004). We identified *Pseudomonas lactucae* CFBP13502 as a potent inhibitor of Xcc8004, mediated by exometabolites specifically induced in the presence of Lysobacterales (formerly Xanthomonadales). Transcriptomic analysis of CFBP13502 revealed upregulation of a gene cluster involved in the biosynthesis of a lipopeptide siderophore biosynthesis. Gene deletion confirmed that this cluster is essential for the growth inhibition of Xcc8004. Furthermore, iron supplementation abolished this inhibitory effect, providing strong evidence for the role of iron chelation. Through comparative metabolomics, we elucidated the structure of a novel family of lipopeptide siderophores, which we named lactuchelins, produced by CFBP13502. Our findings provide the first molecular evidence of competitive exclusion mechanisms at the seed microbiome interface, highlighting lactuchelins as a promising avenue for the development of seed-based biocontrol strategies against seed-borne phytopathogens.

## Introduction

Historically, seeds have been viewed as a means of plant pathogens dispersal which can lead to the emergence of diseases in new geographical areas (Denancé & Grimault, 2022). However, seed transmission is not restricted to plant pathogens as seeds harbor diverse microbial communities (Simonin *et al*., 2022) that can contribute to plant growth and disease suppression (Pal *et al*., 2022; Garin *et al*., 2024). Promoting these beneficial microbes at the expense of plant pathogens through seed inoculation has recently emerged as a promising strategy (Simonin *et al*., 2023; Arnault *et al*., 2024). However, this requires an understanding of the molecular mechanisms involved in intermicrobial competition within this habitat.

Due to their small size and limited nutrient reserves, seeds often host microbial communities dominated by a few key taxa (Newcombe *et al*., 2018; Chesneau *et al*., 2022). Additionally, priority effects are known to lead to nutrient pre-emption for late-arriving microbes (Debray *et al*., 2022) and this has been shown to efficiently explain the establishment of seed inoculums on seedlings (Arnault *et al*., 2024). Consequently, microbial competition might play a decisive role in shaping community composition during seed-to-seedling transmission. Modes of competition between microbes could be exploitative competition where microbes outcompete others by efficiently scavenging scarce resources or interference competition where microbes directly antagonize others (Granato *et al*., 2019). For example, in the case of the seed-borne pathogen *Xanthomonas campestris* pv. *campestris* 8004 (Xcc8004), a strong competitive interaction has been found with *Stenotrophomonas rhizophila*, which ultimately limits Xcc8004 transmission from seed to seedling. This competition is mediated by T6SS interference competition (Garin *et al*., 2024) and potentially exploitative competition (Torres-Cortés *et al*., 2019).

While carbon is often considered the primary limiting nutrient in seeds, micro-elements availability presents also a significant challenge for microbial establishment. For instance, iron plays an important role in *Pseudomonas putida* establishment on seeds (Molina *et al*., 2005). The ability to efficiently acquire iron can thus be a major determinant of microbial success during seed transmission. Iron acquisition is usually achieved through the secretion of siderophores, secondary metabolites that possess different iron-chelating functional groups such as catechols, hydroxamates, carboxylates, or phenolates (Schalk, 2025). Siderophores are well-known for their role in microbe-microbe interactions and competition (Kramer *et al*., 2020).

Pyoverdines are a well-characterized group of siderophores produced by *Pseudomonas* species that contribute to the fluorescence property of these bacteria (Cezard *et al*., 2015). Pyoverdine biosynthesis genes are highly conserved, present in 97% of sequenced *Pseudomonas* genomes (Gu *et al*., 2024). Non-fluorescent pseudomonads can however produced other siderophores such as corrugatin (Risse *et al*., 1998) and its derivatives ornicorrugatin (Matthijs *et al*., 2016) and histicorrugatin (Matthijs *et al*., 2016). These peptidic siderophores are acylated with a lipid chain of variable length. Lipopeptide siderophores are highly prevalent in marine bacteria, since the amphiphilic properties of these molecules could limit siderophore diffusion via membrane attachment (Sandy & Butler, 2009; Gauglitz *et al*., 2014; Robertson *et al*., 2018; Galica *et al*., 2021). However, lipopeptide siderophores are also present in a number of plant-associated bacteria, such as strains of the *Burkholderia cepacia* group (ornibactins, Stephan *et al*., 1993), *Cupriavidus taiwanensis* (taiwachelins, Kreutzer & Nett, 2012), *Herbaspirillum seropedicae* (serobactins, Rosconi *et al*., 2013), *Variovorax boronicumulans* (variochelins, Kurth *et al*., 2016) or *Azotobacter chroococcum* (crochelins, Baars *et al*., 2018).

Despite the importance of iron acquisition in microbial competition, the role of siderophore-mediated interactions in seed-borne microbial communities remains underexplored. By analysing seed-borne bacterial strains that are highly competitive against Xcc8004 during seed-to-seedling transmission (Torres-Cortés *et al*., 2019; Simonin *et al*., 2023), we identified a non-fluorescent strain belonging to the *Pseudomonas lactucae* species (Sawada *et al*., 2021) that significantly reduced Xcc8004 growth. Compounds responsible for this activity are related to a new lipopeptide siderophore family that we propose to name lactuchelins. Lactuchelins production in *P. lactucae* CFBP13502 is enhanced during interactions with specific bacterial strains including Xcc8004. The gene cluster involved in the biosynthesis of this compound is distributed in eight species of *Pseudomonas* including several strains isolated from seed. Investigating how seed-associated bacteria utilize these siderophores to outcompete plant pathogens could provide novel strategies for microbiome-based disease management in crops.

## Methods

### Bacterial strains and culture media

Thirty bacterial strains covering the diversity of seed microbial communities (Simonin *et al*., 2022) were selected (**Table S1**). All these strains were isolated from radish seed samples collected in 2013, 2014 and 2015 (Rezki *et al*., 2018) except one strain isolated from radish flower in 2016 (Chesneau *et al*., 2020). *Xanthomonas campestris* pv. *campestris* 8004 (Xcc8004) is a spontaneous rifampicin-resistant strain (Turner *et al*., 1984), which derives from *X. campestris* pv. *campestris* NCPPB 1145, a strain isolated from cauliflower. All strains were routinely grown in 1/10 strength Tryptic Soy Broth (TSB 10% : 17 g.l^-1^ tryptone, 3 g.l^-1^ soybean peptone, 2.5 g.l^-1^ glucose, 5 g.l^-1^ NaCl and 5 g.l^-1^ K_2_HPO_4_) at 28°C, supplemented when needed with 15 g. l^-1^ of agar and 50 µg.ml^-1^ of rifamycin.

Mono-culture and co-culture of the 30 selected bacterial strains with Xcc8004 were performed at 28°C under constant shaking (150 rpm) during 24h using a starting OD_600_ of 0.02. Culture supernatant was filtered through 0.22 µm filters. The resulting cell-free supernatant (CFS) was mixed 40% (v/v) with 50% (v/v) of 1/5 strength TSB and 10% (v/v) of a fresh cell suspension of Xcc8004 at an initial OD_600_ of 0.2. Three hundred µl of the resulting mix was added per well of 96 well-plates. OD was recorded every hour during 24 hours, under constant shaking (150 rpm) at 28 °C (**Fig. S1**). Iron supplementation was performed by adding a range of FeCl_3_ concentration (from 0.3 µM to 50 µM final concentration) in the coculture or in the CFS. Times of generation were calculated using library growthcurver R package (version 0.3.1). Differences between generation times were assessed by Kruskall-Wallis tests followed by Dunn’s *post-hoc* tests.

### Genome sequencing and annotation

Genome sequences of radish-associated bacterial strains were initially obtained through HiSeq 4000 PE150 sequencing (Torres-Cortés *et al*., 2019). Genomes sequences were assembled with SOAPdenovo 2.04 (Luo *et al*., 2012) and VELVET 1.2.10 (Zerbino & Birney, 2008) and annotated with prokka 1.14.6 (Seemann, 2014).

In addition, DNA of *P. lactucae* 13502 was subjected to long-read sequencing. High-molecular weight (HMW) DNA was extracted using the Masterpure kit (Lucigen Corporation, USA) with slight modifications. A cell pellet was resuspended in PBS to obtain OD_600_ 2.0. 1 mL of OD_600_ 2.0 culture was centrifuged, decanted, then resuspended in 300 uL TE buffer with 2.5 uL lysozyme (5 mg/mL) and 1 uL RNaseA (5 mg/mL). The solution was incubated at 37°C for 45 min. Cells were lysed by adding 300 uL of MasterPure 2x Tissue and Cell Lysis Solution and 1 μL of Proteinase K (50 mg/mL). Samples were incubated at 65°C for 15 min then placed on ice for 5 min. HMW DNA was further purified using phenol-chloroform extraction. Nanopore libraries were created using the SQK-LSK109 kit and sequenced on a MinION (Oxford Nanopore Technologies, UK) with a R9.4.1 flow cell. Basecalling, adapter removal, and barcode removal of the nanopore reads was done with Guppy 4.0 software (Oxford Nanopore Technologies, UK). Illumina libraries were prepared with the Illumina DNA prep kit and sequenced with 2X150bp reads on a NovaSeq 6000 (Illumina, USA). Illumina reads were trimmed and cleaned using BBtools (https://jgi.doe.gov/data-and-tools/bbtools) (Lui *et al*., 2021). Illumina and nanopore reads were used as input to Unicycler (Wick *et al*., 2017) for hybrid genome assembly. Unicycler was run with default parameters. Visualization of the gene cluster was performed with geneviewer 0.1.10 (Ven der Velden N, 2025).

### RNA-Seq experiments

Transcriptome analysis of *P. lactucae* CFBP13502 was performed during co-culture with two strains (*Stenotrophomonas rhizophila* CFBP13503 and Xcc8004) that produced a CFS with bacterial growth inhibition potential. These transcriptomes profiles were compared to co-culture of *P. lactucae* CFBP13502 with two strains (*Pantoea agglomerans* CFBP13505 and *P. viridiflava* CFBP13507) that did not trigger CFS with Xcc8004 inhibitory activity. Four independent replicates were performed per co-culture at a starting OD_600_ = 0.02. Total RNA of each strain was isolated by a modification of the method of (Piper *et al*., 1999). A 2-ml suspension of co-culture after 6 h growth was centrifuged 30 s at 14,000 × g. The pellet was resuspended in 600 μl of hot (65°C) Trizol (Invitrogen, Cergy Pontoise, France) and the mixture was shaken at 65°C for 5 min. After 5 min at room temperature, the cell debris was removed by centrifugation (2,500 × g, 5 min at 15°C) and the supernatant was transferred to a fresh tube. Chloroform (120 μl) was added to 600 μl of the supernatant and the mixture was vortexed vigorously 30 s. After 2 min at room temperature, the phases were separated by centrifugation at 12,000 × g for 15 min at 4°C and the upper phase (300 μl) was removed to a fresh tube. Isopropyl alcohol (250 μl) was added and the mixture was incubated at room temperature for 10 min. The precipitated RNA was collected by centrifugation at 12,000 × g for 20 min at 4°C. The supernatant was removed and the pellet was washed with 500 μl of 75% ethanol. A new pellet of RNA was collected by centrifugation at 7,500 × g for 5 min at 4°C and was air dried to near completion. The RNA was dissolved in 89 μl of sterile double-distilled H2O, and the preparation was treated with 1 μl of DNase (Ambion Applied Biosystems, Courtaboeuf, France) (2 U μl–1) to remove contaminating DNA during 1 h at 37°C. The preparation was heated 10 min at 75°C to inactivate the DNAse and RNA was ethanol precipitated and dissolved in 50 μl of double-distilled H2O.

RNA-Seq libraries were prepared with the QIAseq FastSelect −5S/16S/23S and QIAseq stranded total RNA Lib kits following manufacturer’s instructions (Qiagen, Venlo, Netherlands). RNA fragmentation was performed at 89°C for 6 min. The quality of the resulting libraries was assessed with a Bioanalyzer (Agilent, Santa Clara, CA) and concentration of libraries and subsequent equimolar pool was quantified with quantitative PCR using Illumina primers (Roche, Basel, Switzerland). The equimolar pool was sequenced with Illumina NextSeq550 using High Output 150 cycles cartridge (Illumina, San Diego, CA)

A median of 25 million paired-end reads (min= 16, max = 32) was obtained per sample. Quality and adapter trimming was performed with Trim Galore 0.6.3 (https://github.com/FelixKrueger/TrimGalore) using a Phred score of 20. Reads were mapped on *P. lactucae* CFBP13502 genome sequence with Salmon (Patro *et al*., 2017). Differentially-expressed genes (DEGs) were determined using DeSeq2 v1.36 (Love *et al*., 2014) at an adjusted *p-value* < 0.01 (Wald Test) and a logarithmic fold change (|log_2_FC| > 2) estimated with apeglm v1.18 (Zhu *et al*., 2019). Proteins function were inferred with PaperBLAST (Price & Arkin, 2021). Iron gene repertoire of *P. lactucae* CFBP13502 was estimated with FeGenie (Garber *et al*., 2020).

### Construction of P. lactucae CFBP13502ΔltcJ

Deletion of *ltcJ* (APEGCL_17050) was performed by allelic exchange using the suicide vector pEX18Tc (Hoang *et al*., 1998). The deletion plasmid pEX18Tc-Δ*ltcJ* was constructed using the TEDA cloning procedure (Xia *et al*., 2019). Briefly, pEX18Tc was digested with XbaI (New England Biolabs) followed by shrimp alkaline phosphatase treatment (Phusion High-Fidelity DNA polymerase; New England Biolabs). *ltcJ* flanking regions were PCR-amplified from CFBP13502 with the Phusion High-Fidelity DNA polymerase and the primer pairs listed in **Table S2**. The dephosphorylated pEX18Tc vector and PCR products were purified using the NucleoSpin Gel and PCR Clean-up kit (Macherey-Nagel). TEDA reaction was then carried out by mixing 150 ng of pEX18Tc with the corresponding PCR products at a molar ratio of 1:4. One hundred microlitres of *E. coli* DH5α were transformed with 5 μL of TEDA reaction using the Inoue transformation procedure (Sambrook & Russell, 2006) modified by Xia et al. (2019). Amplicon insertions were validated by colony PCR with the primer pair M13F/M13R. Plasmids (pEX18Tc-Δ*ltcJ*) were extracted with the NucleoSpin plasmid kit (Macherey-Nagel), and insertion regions were verified by sequencing. *E. coli* MFDpir was transformed with pEX18Tc-Δ*ltcJ* using the modified Inoue method. Plasmids were transferred to *P. lactucae* CFBP13502 by conjugation. *P. lactucae* CFBP13502 transconjugants were selected on TSA10 supplemented with tetracycline (20 μg/mL). The resulting colonies were grown in TSB10 (28°C, 120 rpm, 3 h) and bacterial suspensions were spread on TSA10 supplemented with 5% saccharose. Allelic exchanges were validated by PCR and sequencing.

Deletion mutants were complemented in trans with the plasmid pBBR-MCS3. The pBBR-MCS3 plasmid was digested by XhoI and XbaI and ligated with the PCR-amplified product of *ltcJ* obtained from CFBP13502 with the Phusion High-Fidelity DNA polymerase and the primer pairs listed in **Table S2.** Plasmids (pBBR-*ltcJ*) were extracted with the NucleoSpin plasmid kit (Macherey-Nagel), and insertion regions were verified by sequencing. *E. coli* MFDpir (Ferrières *et al*., 2010) was transformed with pBBR-*ltcJ* using the modified Inoue method. Plasmids were transferred to *P. lactucae* CFBP13502 by conjugation. *P. lactucae* CFBP13502 transconjugants were selected on TSA10 supplemented with tetracycline (20 μg/mL). The resulting colonies were spread on TSA10 (28°C, 48h) and the presence of the pBBR-*ltcJ* plasmid was validated by PCR.

### Genome-wide mutant fitness assays

We performed pooled mutant fitness assays as described previously (Wetmore *et al*., 2015; Price *et al*., 2018). The Xcc8004 mutant library (Luneau *et al*., 2022) was revived in MOKA rich medium (Yeast Extract 4 g/l, Casamino acids 8 g/l, K2HPO4 2 g/l, MgSO4.7H2O 0.3 g/l) supplemented with rifampicin, 50 µg/ml; kanamycin: 50 µg/ml; tetracycline: 5 µg/ml. Culture was left to grow at 28°C under constant agitation (150rpm) until reaching mid-log phase. Samples of the culture of Xcc8004 pooled mutants were collected as the “Time0” controls. The remaining culture was pelleted and washed two times then resuspended in 10mM MgSO4.

We incorporated the Xcc8004 mutant library (starting OD_600_ = 0.02) in Xcc8004+13502 cell-free supernatant supplemented with TSB20, and compared, as controls, the fitness of the library in three other cell-free supernatant : the mutant library grown : (i) in PBS, (ii) in Xcc8004 cell-free supernatant and (iii) in 13502 cell-free supernatants (Cultures were performed following 1/4/5 ratio described in “Phenotype screening for Xcc8004 inhibition” section). We collected the mutant population in our various conditions after 24 hours of growth, and extracted genomic DNA from the Time0 and condition samples using NucleoSpin® 96 Food kit (Macherey-Nagel) following the supplier’s recommendations. 200 ng of DNA were amplified using Q5 Polymerase (NEB) and BarSeq primers (Wetmore *et al*., 2015). PCR cycling conditions were as follows: denaturation at 98°C (4 min), 25 cycles at 98°C (30 sec), 55°C (30 sec) and 72°C (30 sec), and a final elongation at 72°C for 5 min. Amplicons were purified with magnetic beads and pooled in equimolar concentration. Concentration of the pool was monitored with quantitative PCR (KAPA Library Quantification Kit, Roche) and sequenced on v3 150 lllumina cartridge (Illumina, San Diego, CA).

To analyze our data, we used a mature pipeline developed to acquire fitness from raw sequencing data. Mutant fitness scores were calculated as the normalized log2 ratio of the abundance of its barcode in the library under our experimental conditions versus the “Time0”. Each mutant is associated with a unique barcode; thus the reads of a certain barcode characterize the abundance of the mutant inside the library. Gene fitness is calculated as the weighted mean of the fitness of each individual mutant. Genes fitness were then normalized to correct variation in copy number along the chromosome, the running median along the chromosome is zero. For each fitness score, we performed a t-like test. Finally, to identify significant Xcc8004 genes in 13502+Xcc8004 CFS compared to our controls, we applied two thresholds |fitness|>1 and |t|>5. To infer the function of Xcc8004 genes based on mutant phenotype, we used the SEED database from the fitness browser (fit.genomics.lbl.gov). We also inferred gene function based on scientific articles published using PaperBLAST.

### Comparative metabolic profiling

To identify the metabolite inhibiting Xcc8004 growth we performed comparative metabolic profiling of Xcc8004+13502 CFS, Xcc8004 CFS, 13502 CFS and TSB CFS. The first step was to de-complexified our CFS. For this purpose, Xcc8004+13502 CFS was freeze-dried and submitted to a liquid/liquid extraction with H_2_O *versus* four different organic solvents, ethyl acetate (AcOEt), butanol (BuOH), dichloromethane (DCM) and methyl *tert*-butyl ether (MtBE). Each extract was evaporated then tested as previously described in “Cell-free supernatant assays on Xcc8004 inhibition” section, by replacing cell-free supernatant by our extracts. BuOH extract was selected due to its inhibitory effect on Xcc8004 growth. Thus, we performed liquid/liquid extraction (H_2_O/BuOH) on the fourth cell-free supernatant and analyzed each fraction *via* UPLC-HRMS/MS at a concentration of 100 µg.mL-1. A sample called “Mix” containing an equivalent mixture of the 4 fractions at 100 µg.mL-1 was also prepared and will be used to align all the chromatograms obtained between them and to check the statistical analysis by PCA. Each sample was injected 5 times in both positive and negative modes.

The analysis was performed using a Waters Acquity UPLC H-Class Series system. Chromatographic separation was carried out on an Acquity HSS T3 column (100 mm x 2.1 mm, 1.8 μm) thermostated at 25 °C. The mobile phase consisted of solvent A (0.1% formic acid in water) and solvent B (0.1% formic acid in acetonitrile), with gradient from 0 to 35% of B in 20 min then 35% to 100% of B in 5 min held for 3 min. The flow rate was set at 300 µL.min-1. The injection volume was 1 μl.

Mass spectrometry was performed on a Waters Xevo G2-XS system. The analyses were performed as two separated methods, one operated in positive electrospray ionization (ESI) mode and one in negative ESI mode. Mass data were acquired in MSe Continuum sensitivity mode with a mass range from *m/z* 50 to 2000 with a scan rate of 2 s.

The tuning parameters (positive and negative) were : capillary voltage 500 V, sampling cone 40 V, cone voltage 80 V, source temperature 120 °C, desolvation temperature 500 °C, desolvation gas flow 100 L.h-1 and cone gas flow 1000 L.h-1. The low collision energy was set at 6 V and the high collision energy was a ramp from 15 to 40 V. The collision pressure of argon was constant at 6.5E-3 mbar. Mass correction was performed with leucine enkephalin solution at 200 pg.mL-1, measured at 30 second intervals throughout each injection, at a flow rate of 5 μL.min-1. The ions monitored were *m/z* 556.2766 and 554.2620 in ESI+ and ESI-, respectively.

The data were then processed with the Progenesis QI software. Potential markers of interest were extracted from S-plots constructed following OPLS-DA, and markers were chosen based on their contribution to the variation and correlation within the data set.

### Culture, extraction, purification and identification of lactuchelins

Bulk extraction of the metabolites was performed on liquid cultures of CFBP13502 for 24 h in 2 L of iron-depleted M9 medium at 28 °C. Subsequently, cells were removed from the supernatant by centrifugation (10 min at 5000 g) and filtration through vacuum filtration system with 0.22 µm filter membrane (TPP, Switzerland).

The resulting CFS was then subjected to freeze-drying (22.5 g), redissolved in 1 L of water complemented with iron chloride and extracted 3 times against BuOH (3 x 1 L). The resulting aqueous (LLW) and BuOH (LLB) fractions were concentrated *in vacuo* at 40 °C.

The LLB fraction (1.1 g) was absorbed on Polygoprep C18 bulk media (50-60 µm) with a ratio media/fraction of 2/1, and transferred into a pre-conditioned HyperSep C18 cartridge (5000 mg; 40-60 µm). A ratio bed weight/fraction of 20/1 was applied. The adsorbed metabolites were then eluted with 50 mL of 3 step solvents : water (SW), water/MeOH (50/50 ; SWM), and MeOH (SM). The resulting fractions were concentrated *in vacuo* at 40 °C. Finally, SWM (482 mg) was injected on a preparative HPLC Shimadzu LC-20AP solvent delivery system coupled to an SPD-UV detector and an FRC-10A fraction collector using LabSolutions software and an Thermo Fisher Scientific Hypersil Gold PFP column (150 x 20 mm; 5 μm; 175 Å). The flow rate was set at 18.9 mL.min^-1^ with a gradient of (A) water + 0.1% of formic acid and (B) MeCN + 0.1% formic acid (20% to 60% B in 10 min, 60% to 100% B in 2 min, then 100% B for 6 min) and UV detection was set at 190 nm. The resulting fractions (F1 to F5) were then concentrated *in vacuo* at 40 °C.

F4 (18.4 mg) was subjected to an iron removal protocol. Ferric iron was removed according to literature protocols (Stephan *et al*., 1993). For the following procedures, all work was carried out in iron-free glassware. F4 fraction (13.6 mg) was dissolved in 80 mL of H_2_O, added to 50 mL of a 8-hydroxyquinoline solution in MeOH (290 mg, 1 mM), and stirred at room temperature for 24 h. The MeOH was then removed *in vacuo* at 40 °C, and the remaining aqueous solution was extracted with CH_2_Cl_2_ (5 × 40 mL). The aqueous layer was mixed with a solution of Ga_2_(SO_4_)_3_ (30 mg, 35 µmol) in sulfuric acid (0.1 N) and the mixture was allowed to mix for 30 min at room temperature. After adjustment of the pH at 7, the solution was freeze-dried.

A final purification of the Ga(III)-F4 fraction (12 mg) was accomplished following the previously established preparative purification method to remove excess of gallium and other impurities.

All NMR spectroscopy experiments (1H, 13C and 2D) of lactuchelin (6.5 mg) were recorded as the Ga(III) adduct in DMSO-*d6* on a Bruker Avance NEO 600 MHz spectrometer equipped with a 5 mm DCH cryoprobe. HRMS fragmentation pattern was recorded during the comparative metabolic profiling.

### In vitro growth inhibition assay against Xanthomonas campestris pv campestris 8004

The Xcc8004 growth inhibition activity of fractions and purified lactuchelin was evaluated by an *in vitro* growth inhibition assay. Fractions and purified lactuchelin were resuspended at 1 mg/mL in a 9:1 (v/v) water/DMSO mixture and mixed with 50% (v/v) of a fresh cell suspension of Xcc8004 at an initial concentration of 1600 CFU/µL in TSB 20%. One hundred µl of the resulting mix was added per well of 96 well-plates. The automated MilliDrop Azurevo (Eurofins, France) system was used to identify the effect of fractions and purified lactuchelin on the growth curve of Xcc8004 grown in TSB 10%. The MilliDrop experiments were done according to the MilliDrop protocol. In brief, the bacterial growth was measured in droplets containing the bacterial suspension by a spectrophotometer at different timepoints. The growth curves were finalized after 30 h to prevent coalescence of the droplets. The bacterial droplets were grown at 28 °C. The 96 well-plates used for the generation of droplets were also incubated in a microplate spectrophotometer at 28°C and the OD was recorded every hour during 24 hours, under constant shaking (150 rpm).

## Results

### Identification of a bacterial cell free supernatant inhibiting Xcc8004 growth

Growth of the phytopathogenic strain *Xanthomonas campestris* pv. *campestris* 8004 (Xcc8004) was evaluated in TSB10 medium supplemented with various cell-free supernatants (CFSs) (**Fig. S1**). These CFSs were derived from (*i*) pure overnight cultures of 30 seed-borne bacterial strains or from (*ii*) co-cultures of each bacterial strain with Xcc8004 (**Fig. 1A** and **Table S1**).

**Fig. 1:**
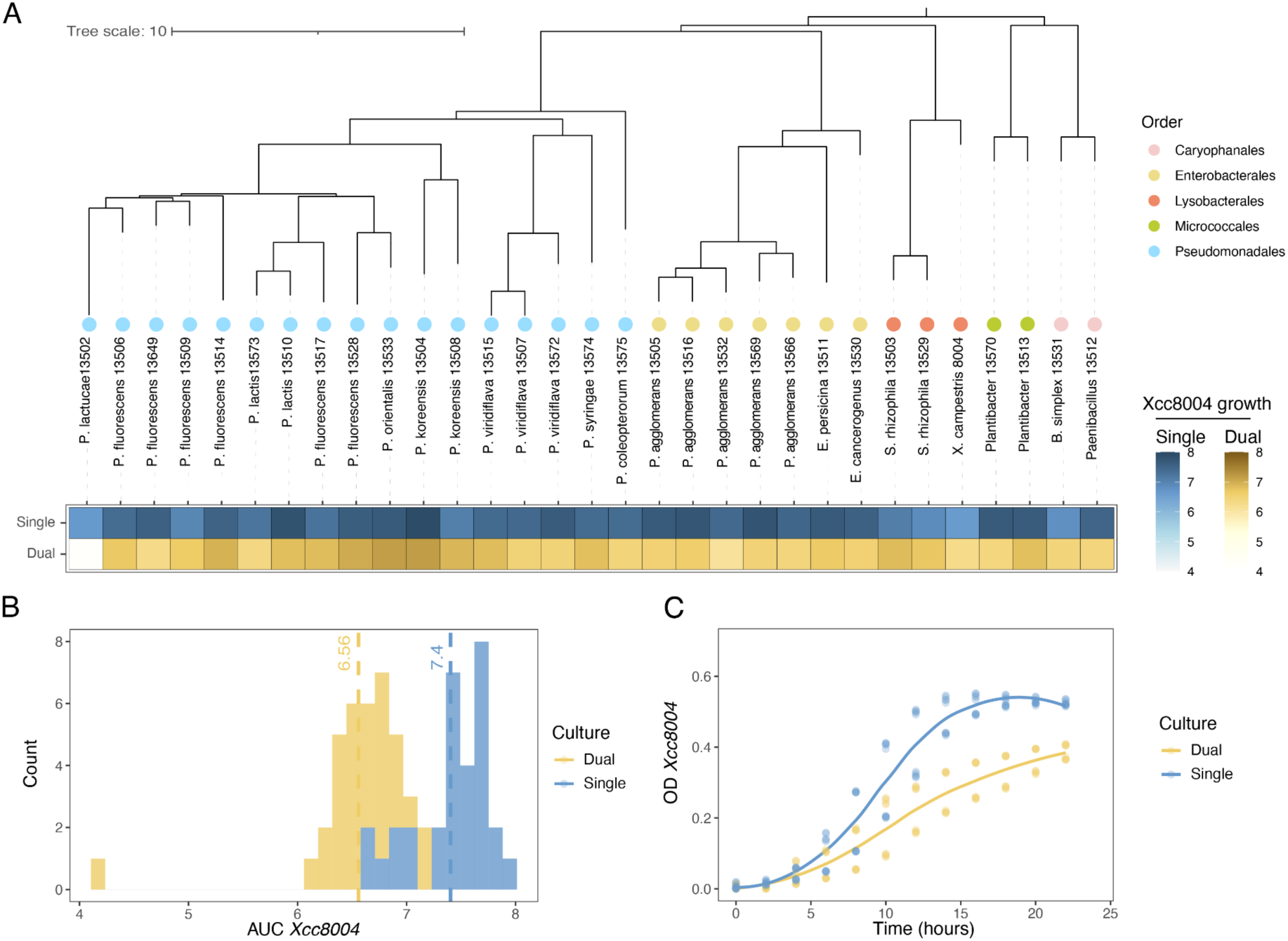
Impact of bacterial cell-free supernatants (CFSs) on *Xanthomonas campestris* pv. *campestris* 8004 (Xcc8004) growth in TSB10. (**A**) Phylogenetic analysis of the 30 seed-borne bacterial strains employed in this work. Heatmap displays Xcc8004 growth (Area under the curve, AUC) following supplementation of CFSs derived from pure cultures of 30 seed-borne bacterial strains (blue-single) or co-culture with Xcc8004 (yellow-dual). (**B**) Distribution of Xcc8004 growth (AUC) after supplementation with CFSs from pure culture (blue-single) or co-culture (yellow-dual). (**C**) Representative growth curves of Xcc8004 supplemented with CFS of a pure culture of CFBP13502 (blue - single) or a co-culture of CFBP13502 and Xcc8004 (yellow - dual).

Most of the CFSs derived from single or co-cultures did not modulate Xcc8004 growth (**Fig. 1B** and **Fig. S2**). The areas under the curve (AUC) of Xcc8004 following supplementation of CFSs collected from single or co-cultures were overall similar, with average AUC values of 7.4 and 6.6, respectively (**Fig. 1B**). However, the CFS collected from the co-culture of *P. lactucae* CFBP13502 and Xcc8004 decreased the AUC of Xcc8004 to 4.2 (**Fig. 1B**), which represented a ∼40% reduction compared to the CFS of *P. lactucae* CFBP13502 alone (**Fig. 1C**). Altogether these results suggest that the co-culture of *P. lactucae* CFBP13502 with Xcc8004 produced metabolite(s) limiting Xcc8004 growth.

To determine whether the growth reduction effect observed was specific to Xcc8004 or could be generalised to other bacterial strains, (*i*) CFS collected from a pure culture of CFBP13502 and (*ii*) CFS collected from the coculture of CFBP13502 and Xcc8004 were applied to cultures of the 30 bacterial strains employed in this study. The growth of 28 strains was inhibited by the CFS derived from the co-culture in comparison with the CFS from the pure culture (**Fig. S3**). However, the magnitude of this inhibitory effect varied, with reductions ranging from 15 to 60% of their AUC. The three strains that were not sensitive to the co-culture supernatant were affiliated to *P. fluorescens* CFBP13506, *P. coleopterorum* CFBP13575 and *P. lactucae* CFBP13502.

### Transcriptome profiling of P. lactucae CFBP13502 highlighted a biosynthetic gene cluster involved in lipopeptide siderophore synthesis

To test whether the observed Xcc8004 growth reductions were also produced by CFBP13502 during interactions with other bacterial strains, CFSs were collected from co-cultures between CFBP13502 and the 30 remaining bacterial strains. A decrease in the AUC of Xcc8004 growth was observed for four of these dual CFSs derived from co-cultures of CFBP13502 with (*i*) Xcc8004, (*ii*) *Stenotrophomonas rhizophila* CFBP13503, (*iii*) *S. rhizophila* CFBP13529 and (*iv*) *Bacillus simplex* CFBP13531 **(Fig. 2A)**.

**Fig. 2:**
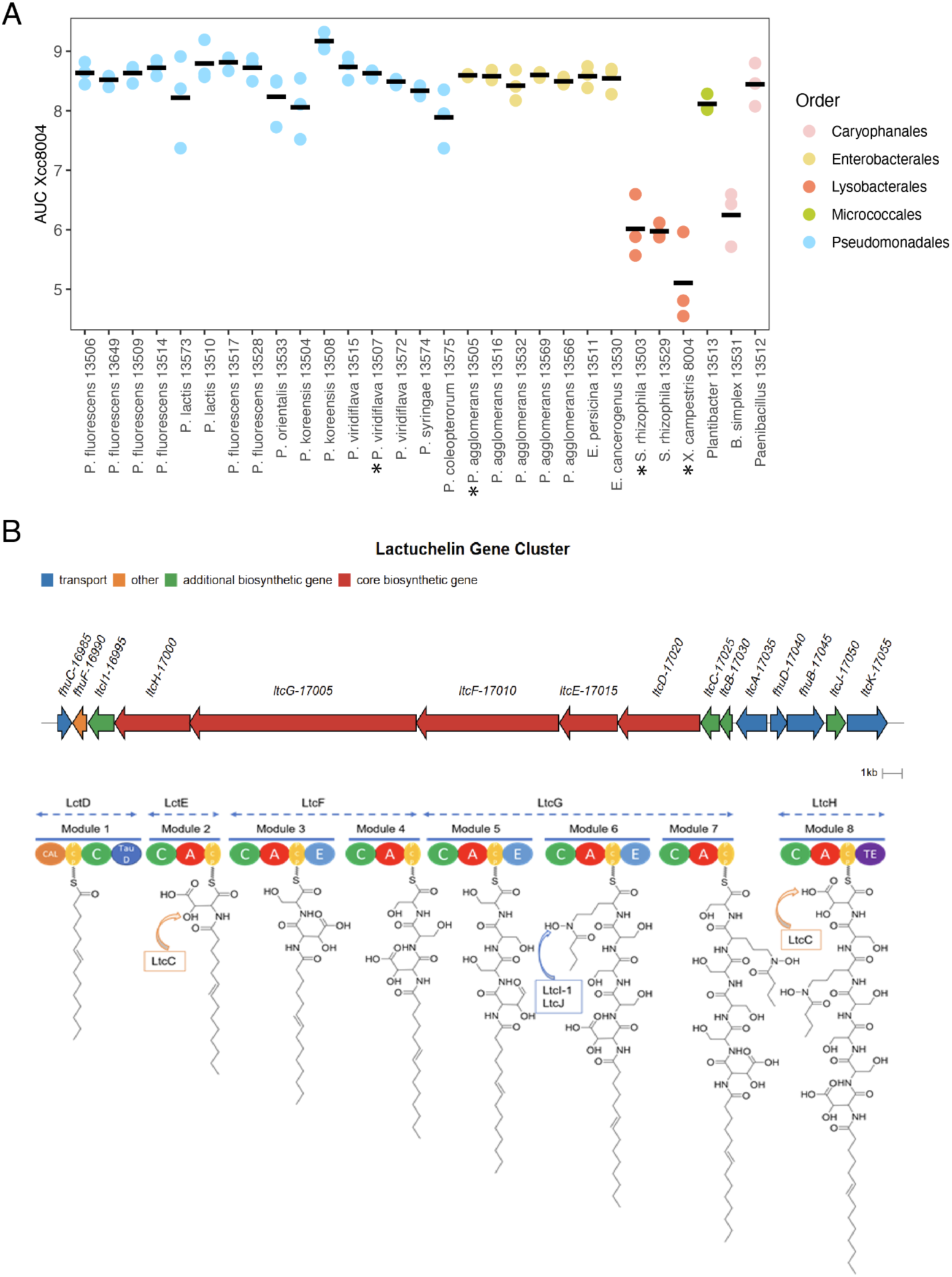
Up-regulation of a biosynthetic gene cluster involved in lipopeptide siderophore production. **(A)** Area under the curve of Xcc8004 growth following supplementation of CFSs derived from co-culture of CFBP13502 and 30 seed-borne bacterial strains. Bacterial strains are colored according to their taxonomic affiliation at the Order level. Stars indicated the cocultures employed for RNASeq experiment. **(B)** Biosynthetic gene cluster involved in synthesis of a lipopeptide siderophore and *in silico* prediction of lactuchelin structure**. (top)** Composition of the biosynthetic gene cluster containing 5 NRPS genes. **(bottom)** Domain architecture of the NRPS proteins and amino acids predicted to be loaded, CAL : Co-enzyme A ligase domain, PCP: Peptidyl-Carrier Protein domain, C: Condensation domain, TauD: hydroxylase, A: Adenylation domain, E: Epimerization domain, TE: Thioesterase domain, C.

Transcriptome profiling of *P. lactucae* CFBP13502 was performed in two co-cultures that produced CFSs with Xcc8004 growth inhibition (*P. lactucae* CFBP13502 with *S. rhizophila* CFBP13503 or Xcc8004) and two co-cultures without reduction of Xcc8004 AUC (*P. lactucae* CFBP13502 with *Pantoea agglomerans* CFBP13505 or *P. viridiflava* CFBP13507, **Fig. 2A**). RNAs were collected after 6h of co-culture, which corresponded to the late exponential growth phase of *P. lactucae* CFBP13502.

Expression was detected in 97% of the 5,687 predicted genes of CFBP13502. Of the 99 differentially expressed genes (DEGs, *P* < 0.01, |log_2_FC| > 2), 29 were associated with iron acquisition and iron gene regulation (**Table S3**). Among these 29 up-regulated genes, 12 were located within a biosynthetic gene cluster (BGC) over a 50 kb region (**Fig. 2B**). According to antiSMASH v7.0 (Blin *et al*., 2023), this BGC possesses sequence similarities to BGC associated with lipopeptide siderophores. The BGC of CFBP13502 is composed of five non-ribosomal peptide synthase (*ltcD-H*) involved in the potential incorporation of the acyl chain and seven amino acids (**Fig. 2B**). Two of these amino acids, aspartate and ornithine, are likely modified by an aspartate β-hydoxylase (*ltcC*, Reitz et al., 2019), a L-ornithine N(5)-monooxygenase (*ltcI_1*) and a N-hydroxyornithine acetyltransferase (*ltcJ*). The BGC also encoded a thioesterase (*ltcB*) and an ABC transporter (*ltcA*) putatively involved in translocation across the cytoplasmic membrane into the periplasm. The uptake of the ferrisiderophore in the perisplasm is probably mediated through a specific TonB-dependent transporter (*ltcK*) and translocation across the inner membrane is likely performed through the ABC transporter FhuBCD. Iron removal is then probably performed through FhuF. Of note *fhuC*, *fhuD* and *fhuB* were the only genes not differentially expressed under our experimental conditions. Moreover other genes potentially involved in siderophore production were located outside the BGC. This is the case for a a small MbtH like protein (APEGCL_19865 (Drake *et al*., 2007)), and a protein-coding gene *ltcI_2* (APEGCL_18425) sharing 72% of identity at the nucleic acid level with *ltcI_1*. According to these genomic predictions, the BGC of *P. lactucae* CFBP13502 is therefore potentially involved in the production of a lipopeptide siderophore. We propose to name this siderophore lactuchelin.

The distribution of the lactuchelin BGC was searched in the 15,013 *Pseudomonas* genomic sequences available at the time of analysis (April 2023). According to protein similarity search this gene cluster is present in 38 *Pseudomonas* strains (**Table S4**). These strains are affiliated to 8 bacterial species (according to the number of shared 15-mers (Briand *et al*., 2021)), which belonged to *P. fluorescens*, *P. putida* and *P. syringae* groups.

### Lactuchelin is responsible for the anti-Xcc8004 activity

In *P. aeruginosa*, production of type II pyoverdine is abolished following deletion of *pvdY2* (Lamont *et al*., 2006) an ortholog of *ltcJ*. A deletion mutant of *ltcJ* was performed in *P. lactucae* CFBP13502 and complemented *in trans* with a complete copy of this gene in the plasmid pBBR-MCS3 (see Methods). The three strains we co-cultured overnight with Xcc8004: CFBP13502 wt, CFBP13502Δ*ltcJ* and CFBP13502Δ*ltcJ-*pBBR-MCS3:*ltcJ*. The corresponding CFSs of these three co-cultures were added in TSB10% medium and the growth of Xcc8004 was monitored. A CFS collected from a pure culture of CBP13502 was also added in TSB10% as a control. We observed a significant (*P*< 0.05) reduction of Xcc8004 growth (**Fig. 3A**) in media supplemented with the CFS collected from the co-culture CFBP13502 wt and Xcc8004 in comparison to the CFS of a pure culture of CFBP13502 wt. In contrast, no significant difference in Xcc8004 growth was observed following supplementation of the CFS derived from CFBP13502Δ*ltcJ a*nd Xcc8004 (**Fig. 3A**). Complementation of CFBP13502Δ*ltcJ* with pBBR-MCS3:*ltcJ* partially restored the growth inhibition of Xcc8004. Altogether these results indicated that the BGC of CFBP13502 was responsible for the growth limitation of Xcc8004 observed following supplementation of the CFS.

**Fig. 3:**
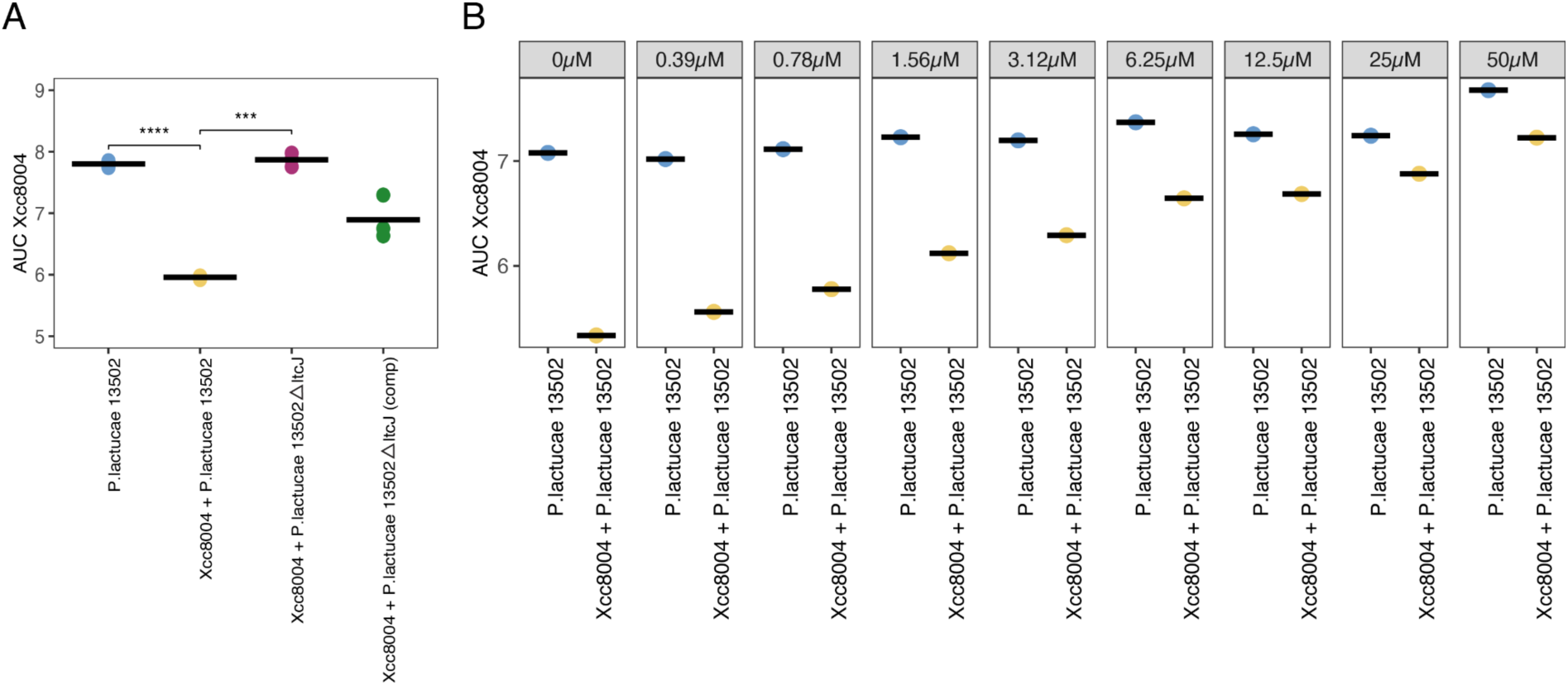
**(A) *ltcJ* mutation abolished the anti-Xcc8004 activity.** Xcc8004 growth was monitored in TSB10 medium supplemented with CFSs from CFBP13502 (yellow) and co-cultures of Xcc8004 and CFBP13502 (blue), CFBP13502Δ*ltcJ* (purple) and CFBP13502Δ*ltcJ* pBBRMCS3-*ltcJ* (green). **(B) Xcc8004 growth inhibition is relieved by FeCl_3_ supplementation.** Xcc8004 growth was monitored in TSB10 medium supplemented with CFSs from CFBP13502 (blue) and co-culture of Xcc8004 and CFBP13502 (yellow). Each CFS was supplemented with various concentrations of FeCl_3_ ranging from 0.39µM to 50µM.

If iron-chelation by lactuchelin is responsible for the growth reduction of Xcc8004, supplementation of CFS with FeCl_3_ should alleviate the growth inhibition. A gradual increase in Xcc8004 growth was observed following supplementation of CFS from Xcc8004 and CFBP13502 co-culture with a range of FeCl_3_ concentration (from 0,39µM to 50µM, **Fig. 3B**). These results suggest that iron chelation by lactuchelin produced by CFBP13502 is likely involved in the decrease of Xcc8004 generation time.

To explore the impact of the inhibitory CFS on the physiology of Xcc8004 we employed a library of barcoded transposon mutants (Luneau *et al*., 2022), specifically a method called random bar code transposon site sequencing (RB-TnSeq). The abundances of each barcoded mutant was estimated after growth in TSB10 supplemented with (*i*) PBS, (*ii*) CFS of Xcc8004 and (*iii*) CFS of Xcc8004-CFBP13502. When employing PBS and CFS of Xcc8004 samples as a control group, we identified seven genes with negative fitness values (**Fig. 4A-B**). However, only one genes (Xcc-8004.2406.1 or *mntH)* that encodes a manganese exporter protein was specifically altered (T-test<0.01) in the CFS of Xcc8004-CFBP13502 in comparison to PBS, and CFS from pure cultures of Xcc8004 (**Fig. 4B**).

**Fig. 4:**
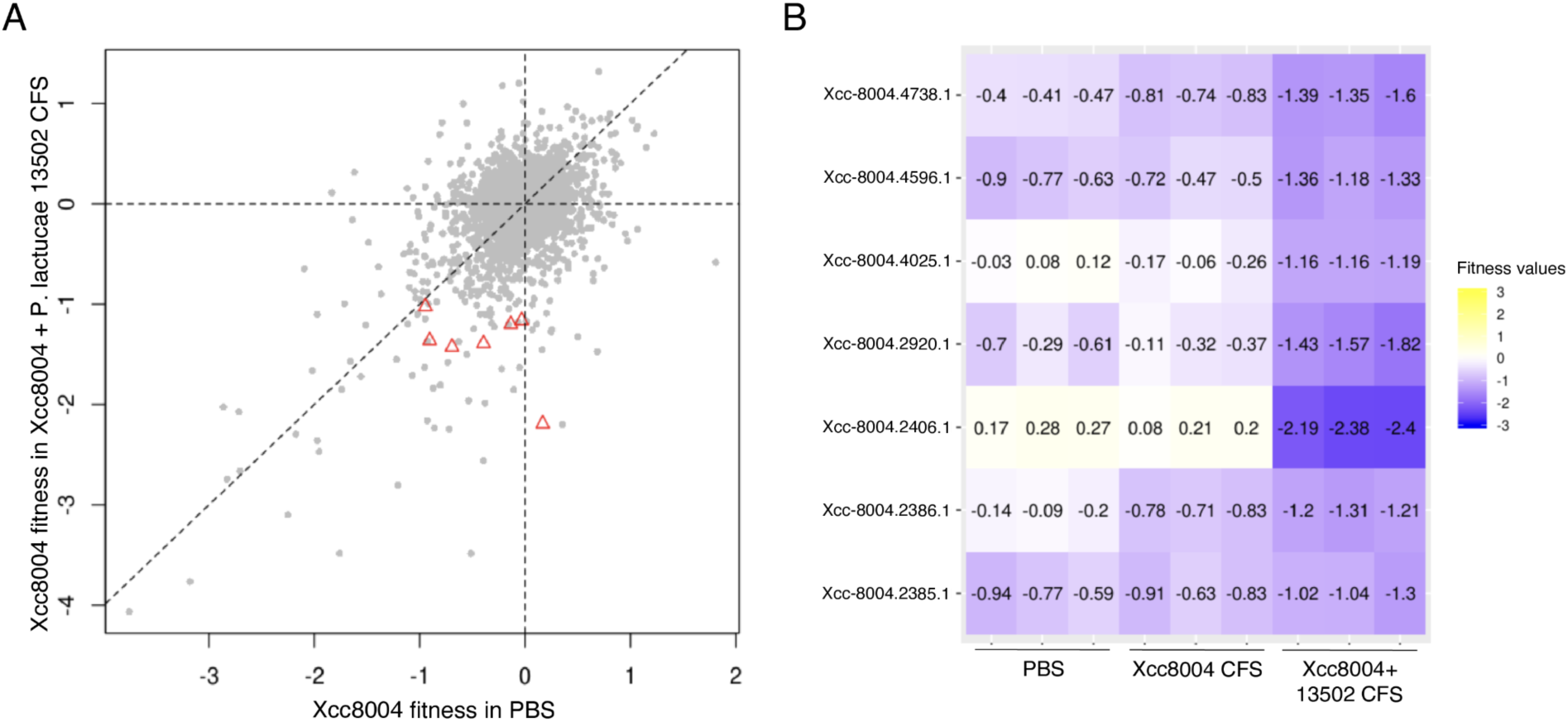
RB-TnSeq screening of *Xanthomonas campestris* pv. *campestris* fitness in inhibitory CFS. **(A)** Comparison of gene contribution to fitness in TSB10 supplemented with PBS (Control) and in TSB10 supplements with CFBP13502 and Xcc8004 CFS. Genes with no significant fitness are in gray. Genes found as differentially represented in CFS condition are shown as red triangles. Each dot corresponds to the mean of 3 replicates. **(B)** Heatmap highlights fitness (gradient color) of the significant Xcc8004 genes in the different conditions.

### Identification of lactuchelins in CFS

To identify the putative lipopeptide siderophore, we performed comparative metabolic profiling of CFSs derived from pure cultures of CFBP13502 and Xcc8004 and CFS from the co-culture of CFBP13502 and Xcc8004. Four different organic solvents (AcOEt, BuOH, DCM and MtBE) were employed for liquid/liquid extraction with H_2_O. The BuOH-extracted fraction had the same anti-Xcc8004 activity than CFS of CFBP13502 and Xcc8004 (**Fig. S4**).

Discriminant analysis of the different conditions in negative ionization mode highlighted 14 compounds enriched in the co-culture of CFBP13502 and Xcc8004 (**Table 1**), potentially involved in bioactivity. Each of them harbors a neutral mass in accordance with a lipopeptide generated by the BGC previously identified (minimal molecular mass 795.64 according to the putative structure of the peptide backbone biosynthesised by the NRPS identified).

**Table 1:** List of metabolites enriched in CFS of coculture of CFBP13502 and Xcc8004, detected in negative mode.

| Peak number | Retention time (min) | m/z | Neutral mass | Anova (p) | Max fold change |
| --- | --- | --- | --- | --- | --- |
| 1 | 19.16 | 515.6610 | 1033.3366 | 2.05E-11 | 11 |
| 2 | 19.23 | 501.1806 | 1004.3758 | 1.60E-09 | 103 |
| 3 | 20.56 | 1023.3334 | 1024.3407 | 2.22E-16 | 36 |
| 4 | 23.08 | 548.1783 | 1098.3712 | 3.66E-15 | 19 |
| 5 | 23.08 | 1051.3659 | 1052.3732 | 4.44E-16 | 31 |
| 6 | 23.34 | 542.6866 | 1087.3878 | 1.28E-13 | 8 |
| 7 | 23.58 | 1121.3526 | 1122.3599 | 1.12E-10 | 13 |
| 8 | 23.70 | 1077.3797 | 1078.3870 | 1.11E-16 | 49 |
| 9 | 23.90 | 529.4484 | 1060.9114 | 1.08E-12 | 40290 |
| 10 | 23.90 | 548.1962 | 1098.4083 | 1.75E-08 | 12 |
| 11 | 24.23 | 1088.3928 | 1089.4001 | 2.07E-08 | 10 |
| 12 | 24.41 | 557.1781 | 1116.3708 | 1.59E-12 | 25 |
| 13 | 24.41 | 562.1927 | 1126.4000 | 5.90E-14 | 32 |
| 14 | 24.41 | 1079.3940 | 1080.4013 | 1.67E-15 | 28 |

*P. lactucae* CFBP13502 was then cultivated in an iron-depleted M9 minimal medium to improve the production of compounds involved in iron chelation. The production of the metabolites described in Table 1 was increased (**Fig. S5**) as well as the inhibitory activity of the CFS (**Fig. S6**). Twenty-six compounds with *m/z* ranging from 951.3821 to 1116.4193 were detected in HRMS analysis in negative mode. They were classified into three series, where mass differences of 2 or 26 between two molecules correspond to an additional unsaturation or a methyl group, respectively. A Δm/z of 52.9114 was calculated for certain compound pairs between series 1 and 2, indicative of a ferric complex [M - 3H + Fe³⁺], while a Δm/z of 23.9581 was observed between compounds of series 1 and 3, corresponding to an aluminum complex [M - 3H + Al³⁺]. Altogether, these findings indicate that series 1 corresponds to the free forms of the siderophores, series 2 to their ferric complexes, and series 3 to their aluminum complexes. Thus, nine siderophores were identified in their free, Fe(III)-bound, or Al(III)-bound forms, differing primarily by the length of their fatty acid chain, ranging from C8 to C16 (**Table S5**).

The inhibitory activity of the different fractions obtained during the purification process was evaluated against Xcc8004 (**Fig. 5**), showing an increase in inhibition with the concentration of the compounds. The fraction F4, with a strong inhibitory activity, contained a purified compound (**1**). The LC-MS/MS analysis in negative mode of this compound revealed an iron-complex form with m/z at 1086.4011, corresponding to [M − 3H + Fe^3+^]- of the Fe(III)-(**1**), which was consistent with a molecular formula of C_43_H_67_FeN_8_O_21_, the associated free form detected in non-supplemented CFS harbour a m/z at 1033.4591 [M-H]-, in accordance with a molecular formula of C_43_H_70_N_8_O_21_. The mass fragmentation pattern in positive mode of the isolated compound (**1**) confirmed the partial amino acid sequence: [fatty acid]-[ꞵOHAsp]-[Ser]-[Ser]- [Ser] (**Fig. S7**). Additional fragments of m/z 417.16 and 286.14, corresponding to a neutral lost of 131.02 validated the presence of a second ꞵ-OH-Asp moiety. Iron complexed with compound (**1**) was removed by treatment with 8-hydroxyquinoline and the free form of compound (**1**) was complexed with gallium for NMR analysis.

**Fig. 5:**
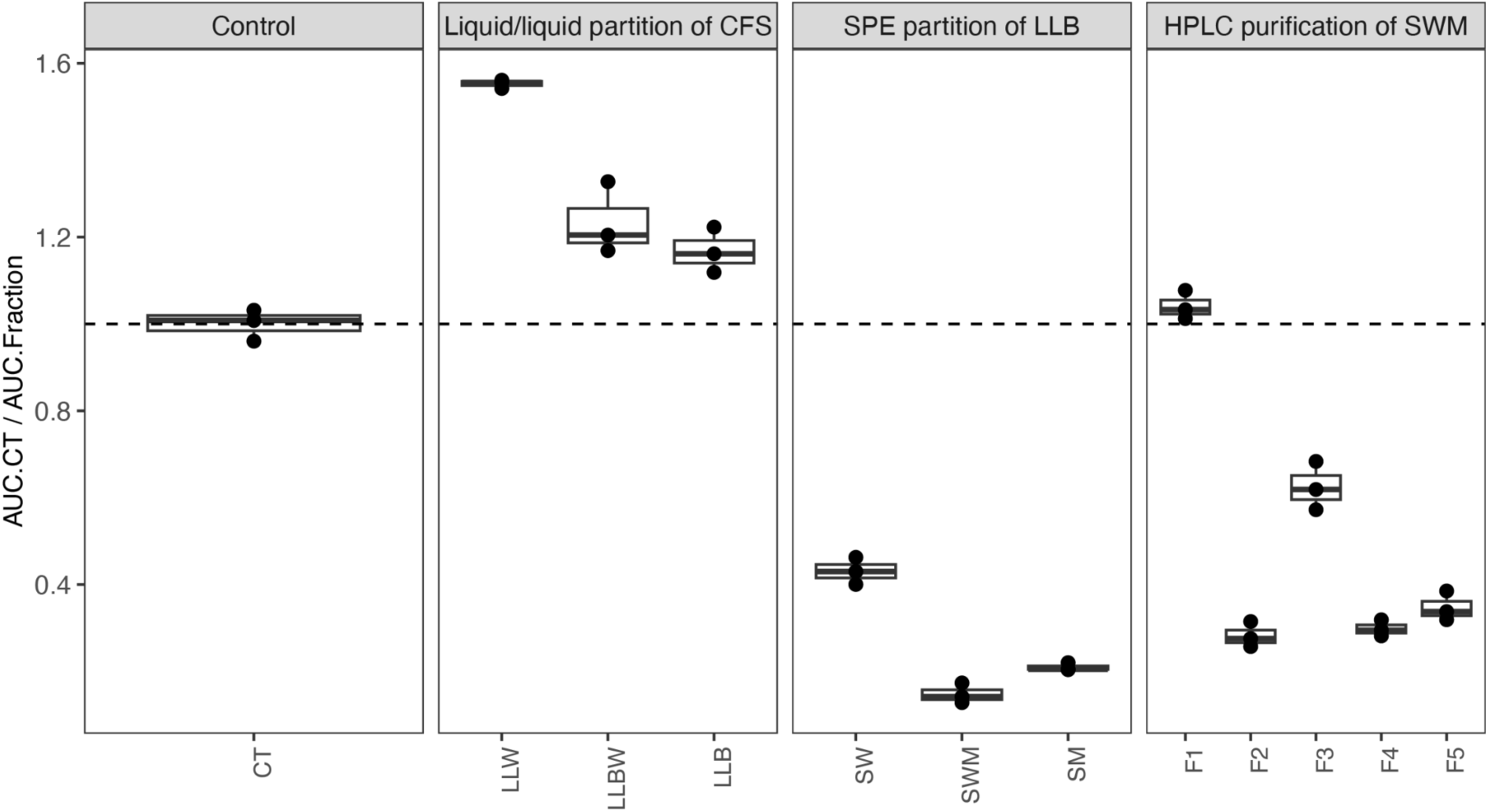
Xcc8004 growth inhibition of the fractions during the purification process of compound 1. Xcc8004 growth was monitored in TSB10 medium supplemented with fractions from CFS of culture of CFBP13502 in iron-depleted M9. Growth inhibition was evaluated by the area under the curve of growth curves (auc). CT: control, LLW: water extract from liquid/liquid partition, LLBW: BOH/water emulsion from liquid/liquid partition, LLB: BuOH extract from liquid/liquid partition, SW: water fraction from SPE partition of BuOH extract, SWM: 50/50 water/MeOH (v/v) fraction from SPE partition of BuOH extract, SM: MeOH fraction from SPE partition of BuOH extract, F1, F2, F3, F4 and F5: fractions obtained from purification of SWM on preparative HPLC.

Structural characterization of Ga(III)-(**1**) was achieved using ^1^H, ^13^C, COSY, HSQC and HMBC experiments (**Table S6** and **Fig. 6**), as well as genetic predictions and MS/MS data. Initial analysis of the ^1^H and COSY spectrum revealed the presence of a lipopeptide containing eight spin systems. This included the four canonical amino acid Ser and three non-canonical amino acids, two ꞵ-OH-Asp and one ornithine (Orn). A final spin system was identified as a 3-hydroxybutanoic acid (Hbu). A tetradecenoic acid moiety was identified by ^1^H and ^13^C NMR and confirmed by MS/MS analysis. HMBC and COSY correlations from the amide protons to adjacent carbonyl carbons and alpha-protons identified the linear peptide sequence as [ꞵOHAsp]-[Ser]-[Ser]-[Ser]-[Orn]-[Ser]-[ꞵOHAsp]. Due to the absence of correlations between Hbu moiety and other spins systems especially between H-5 of the Hbu moiety and the ɑ-carbonyl C-3 of ꞵOHAsp and carbonyl C-7 of Hbu moiety with the H-8 of Orn, the cyclization of the peptide core was confirmed by comparison with the literature data in particular with imaqobactin (Robertson *et al*., 2018). This confirmed the presence of a cyclic depsipeptide linked between Orn and ꞵOHAsp residues via Hbu. Finally, an HMBC correlation between NH-36 and C-37 of the tetradecenoic acid moiety placed it on the NH-36 position coupled via an amide linkage. The location of unsaturation was determined by the analysis of all HMBC and COSY correlations. Unfortunately, the geometry of unsaturation was impossible to resolve due to the impossibility to distinguish H-43 and H-44 both to δH 5.32. The isolated compound (**1**) was named lactuchelin A.

**Fig. 6:**
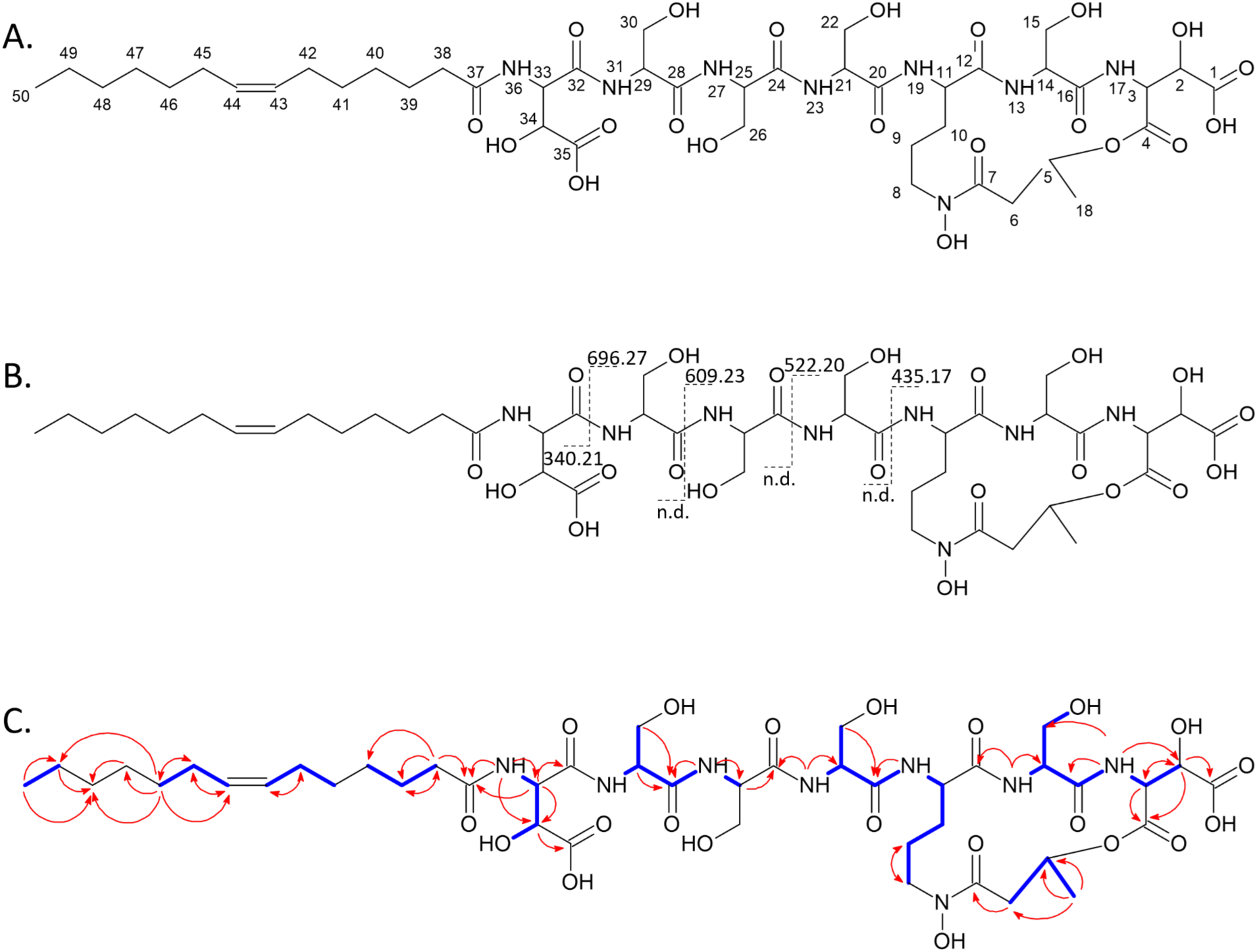
Structural characterization of lactuchelin A. **(A)** Structure of lactuchelin A. **(B)** Fragmentation pattern of lactuchelin A in MS/MS experiment, dashed lines represent the characteristic “y” and “b” ions. **(C)** Key ^1^H−^1^H COSY (blue line) and ^1^H-^13^C HMBC (red arrow) correlations of lactuchelin A.

## Discussion

By screening a collection of seed-borne bacterial strains for antagonistic properties towards the seed-transmitted pathogen *Xanthomonas campestris* pv. *campestris* 8004, we identified a cell free supernatant (CFS) that decreased Xcc8004 generation time of approximately 60%. The antagonistic activity of this CFS was observed during co-culture of the producer strain *P. lactucae* CFBP13502 with four specific bacterial strains tested in this work including Xcc8004. The compounds involved in the growth inhibition of Xcc8004 are a new class of lipopeptide siderophore that we named lactuchelins.

Lactuchelins are newly described hydroxamate siderophores featuring a cyclic depsipeptide core composed of serine, β-hydroxy-aspartate, and N-hydroxyornithine, linked *via* ester and amide bonds, with an additional 3-hydroxybutanoic acid moiety. This type of cyclization has only been reported so far in imaqobactin (Robertson *et al*., 2018). According to metabolomics we were able to identify nine compounds, detected in three different forms: free, complexed with iron or complexed with aluminium. Due to low abundance of these compounds, only one - lactuchelin A - could be fully characterized. The production of lipopeptide siderophores in suites differing by the fatty acid tails is a well-known process, primarily observed in marine environments (Butler, 2005). However, terrestrial examples have also been reported, such as the cupriachelins from the bioplastic producer *Cupriavidus necator* (Kreutzer & Nett, 2012) and serobactins from the grass endophyte *Herbaspirillum seropedicae* (Rosconi *et al*., 2013). The combination of seven amino acid cores with a long fatty acyl chain results in amphiphilic compounds excreted in microbial supernatants. The variation of fatty acid chain length (ranging from C8 to C16) within the lactuchelins series enhances the partitioning of these siderophores into cell membranes. This structural diversity likely improves not only iron uptake (Sandy & Butler, 2009) but also the acquisition of other trivalent metals, such as aluminium (III), in *P. lactucae*, as evidenced by the detection of Al(III) complexes.

Although HRMS analysis indicated that lactuchelins can complex ferric iron and aluminium, several lines of evidence indicated that chelation of iron is involved in the growth inhibition of Xcc8004. First FeCl_3_ supplementation of the CFS completely relieved growth inhibition. Second, RB-TnSeq experiments have highlighted a decrease of fitness of *mntH* deficient mutants in Xcc8004 following supplementation with CFS from the coculture of Xcc8004 and CFBP13502. In *Neisseria meningitidis*, *mntH* encodes a manganese exporter protein that regulates the intracellular ratio of Mn/Fe conferring protection against manganese toxicity at low iron concentration (Veyrier *et al*., 2011). Other *Xanthomonas* species are sensitive to iron deprivation by siderophores. For instance pyoverdine productions by *P. putida* KT2440 can inhibit the growth of *Xanthomonas fragariae* (Henry *et al*., 2016). Finally, the growth inhibition of a large number of strains (n=27) following CFS supplementation of the co-culture of Xcc8004 and CFBP13502 supports the chelation of an essential element for bacterial growth. The three strains that were not impacted by this CFS corresponded to *P. lactucae* CFBP13502, *P. coleopterorum* CFBP13575 and *P. fluorescens* CFBP13506. The two first strains (CFBP13502, CFBP13575) possessed the BGC of lactuchelins that include the TonB-dependent transporter LtcK potentially involved in lactuchelins uptake. While *P. fluorescens* CFBP13506 does not have this BGC, an ortholog of LtcK is encoded in its genome sequence. Therefore, this strain can potentially utilize lactuchelins produced by *P. lactucae* CFBP13502 *via* xenosiderophore utilization, a strategy called siderophore piracy (Traxler *et al*., 2012).

In comparison to pyoverdine, the distribution of lactuchelin biosynthesis genes within *Pseudomonas* is much less important, representing 0.25% of the genome sequences availables versus 97% for pyoverdine (Gu *et al*., 2024). Lactuchelins BGC is found in all strains of *P. lactucae* (*P. fluorescens* group, (Sawada *et al*., 2021), *P. californiensis* (*P. syringae* group, (Carvalho *et al*., 2022) and *P. coleopterorum* (*P. putida* group, (Menéndez *et al*., 2015) species. Interestingly these lactuchelin-producing strains do not possess pyoverdine BGC. This “mutual exclusion” between siderophores is not frequent. Indeed, most pyoverdine-producers strains can usually synthesize other siderophores (known as secondary siderophores because of their lower affinity for ferric iron) including the lipopeptide siderophores ornicorrugatin (Matthijs *et al*., 2008) and histicorrugatin (Matthijs *et al*., 2016). The absence of pyoverdine BGC in lactuchelin-producers is probably due to the fact that producing these two siderophores is too costly to maintain. Lactuchelin production could reflect an iron acquisition strategy that is not solely based on the diffusion of siderophores in the external environment (as is the case with pyoverdines). Indeed the amphiphilic nature of lactuchelin could limit siderophore diffusion through membrane association (Gauglitz *et al*., 2014). As a result, moderate siderophore secretion could then limit their use by individuals other than the producers (Kümmerli, 2023).

The induction of lactuchelin production by *P. lactucae* CFBP 13502 is specifically triggered during coculture with three *Lysobacteraceae* strains (Xcc8004, *Stenotrophomonas rhizophila* CFBP13503 and CFBP13529) and one strain of *Bacillus simplex* (CFBP13531). This suggests that iron concentration is not the primary driver of lactuchelin biosynthesis under our experimental conditions. Rather, this induction may result from signaling molecules that stimulate lactuchelin biosynthesis. Previous studies have demonstrated that siderophore production can be regulated by various factors, such as quorum sensing (Popat *et al*., 2017), ions (Złoch *et al*., 2016), and primary metabolites. For example, histidine represses the histidine utilization repressor HutC, thereby inducing pyoverdine synthesis in *P. fluorescens* SBW25, independent of iron limitation (Naren & Zhang, 2020). Further investigation of the interactions between Xcc8004 and *P. lactucae* 13502 is necessary to better understand the mechanisms driving its induction.

In conclusion, *P. lactucaea* CFBP 13502, through its production of the newly identified lipopeptide siderophore lactuchelin, emerges as a promising candidate for targeted microbial seed treatments to reduce the impact of the seed-borne pathogen Xcc8004. While lactuchelin exhibits broad antimicrobial activity within the seed microbiota, its specific induction by *Lysobacteraceae* may facilitate a precise and efficient biocontrol strategy, minimizing unintended disruptions to native microbial communities and the surrounding environment.

## Acknowledgements

This work was supported by the French National Research Agency (ANR-17-CE20-0009-01 and ANR-20-PCPA-0009) and funding from the France-Berkeley Fund (Engineering a pathogen-resistant seed microbiome). We thank Adam Deutschbauer and Trenton Owens for advice about mutant library experiments as well as the ANAN platform (SFR QuaSaV) for RNA sequencing.

## Supplementary figures

**Fig. S1:**
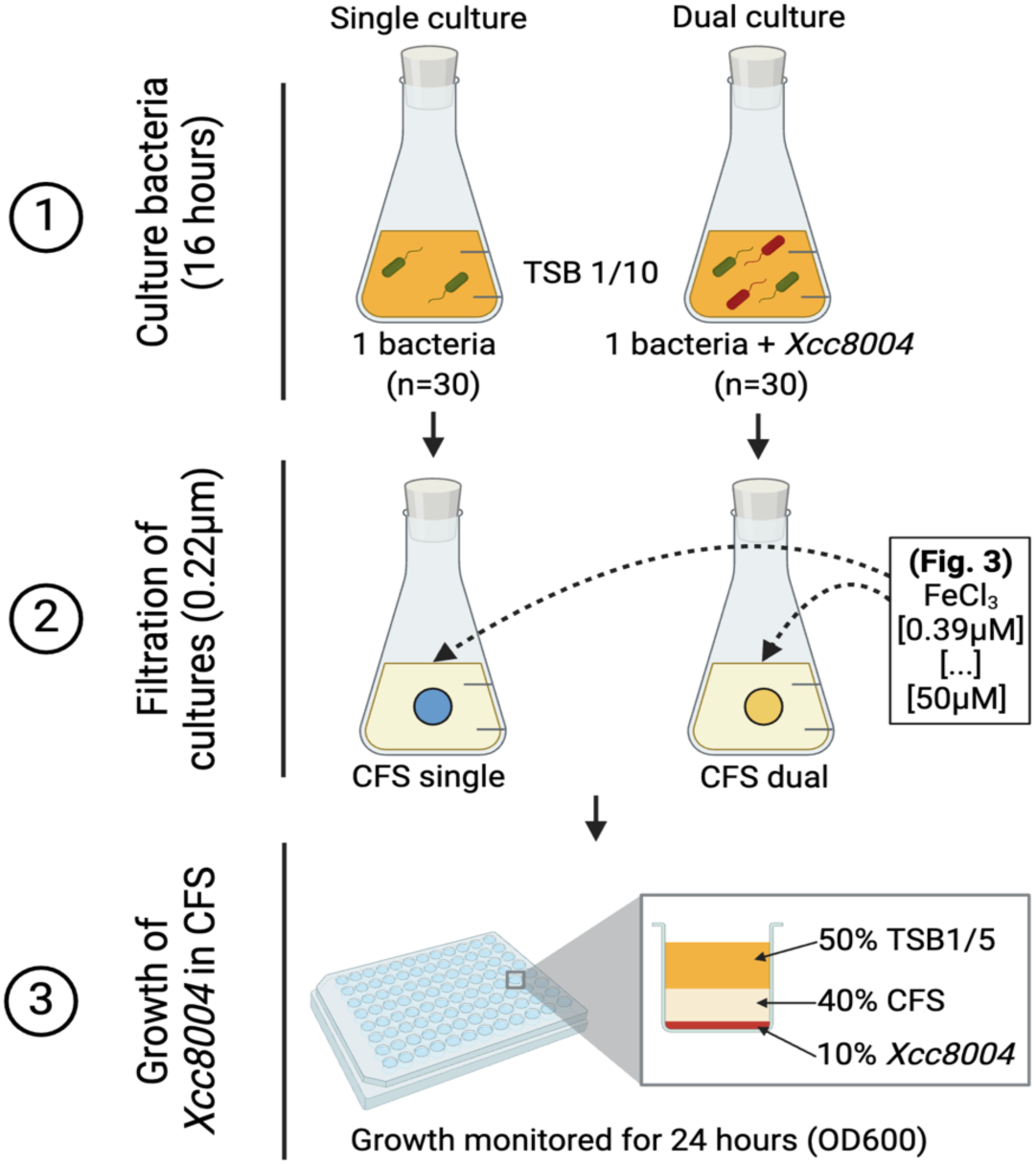
FlowChart culture-free supernatant experiments (CFSs)

**Fig. S2:**
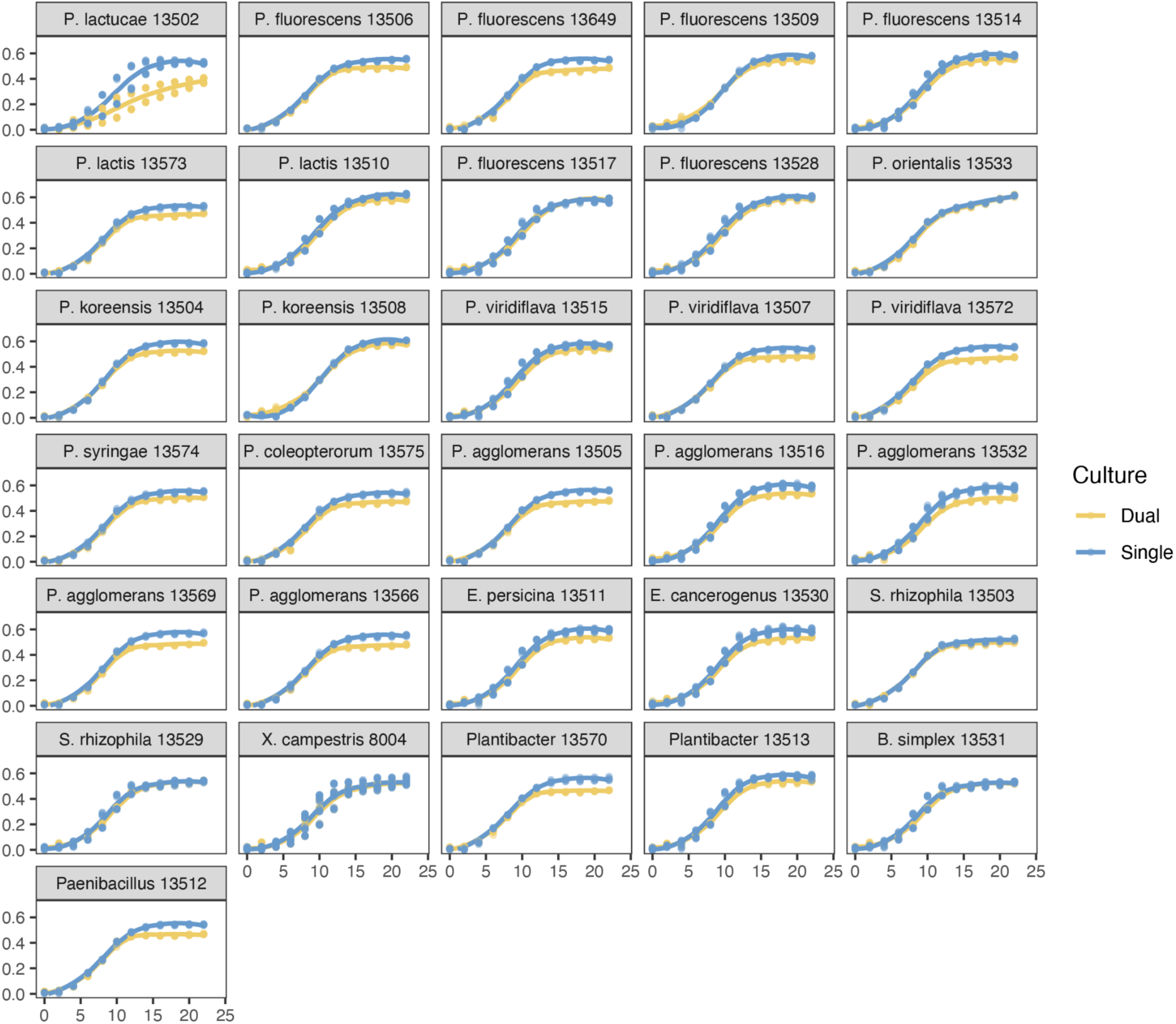
Xcc8004 growth following supplementation with CFSs. Xcc8004 growth was monitored in TSB10 medium supplemented with single or dual cell-free supernatants (CFSs).

**Fig. S3:**
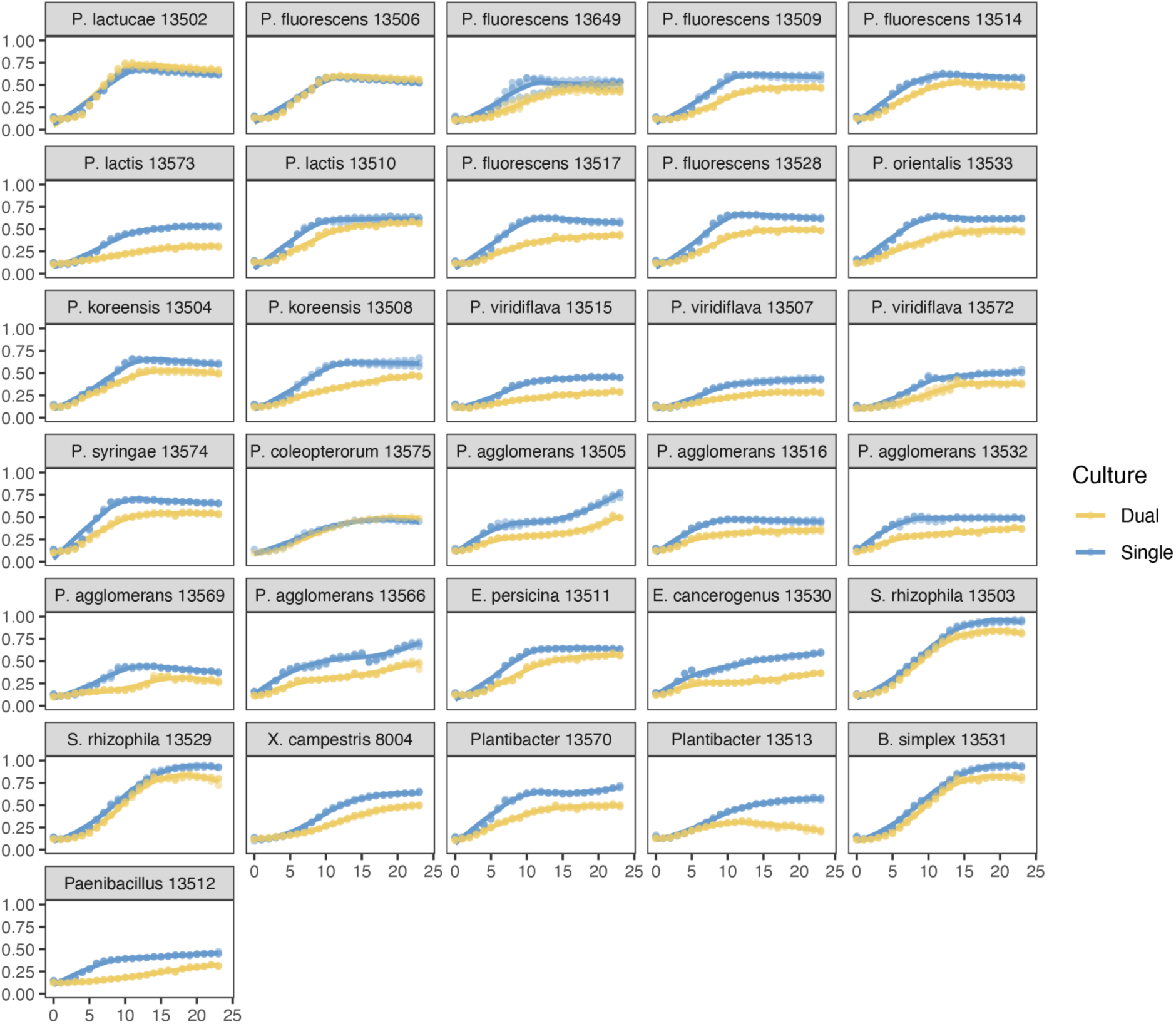
The CFS of CFBP13502 and Xcc8004 inhibited the growth of a wide range of bacterial taxa. 31 bacterial strains growth (panels) were measured for 24 hours in TSB10% supplemented with CFS of CFBP13502 (single - blue) or CFS of CFBP13502+Xcc8004 (dual - yellow), using spectrophotometer. Three replicates were performed for each condition.

**Fig. S4:**
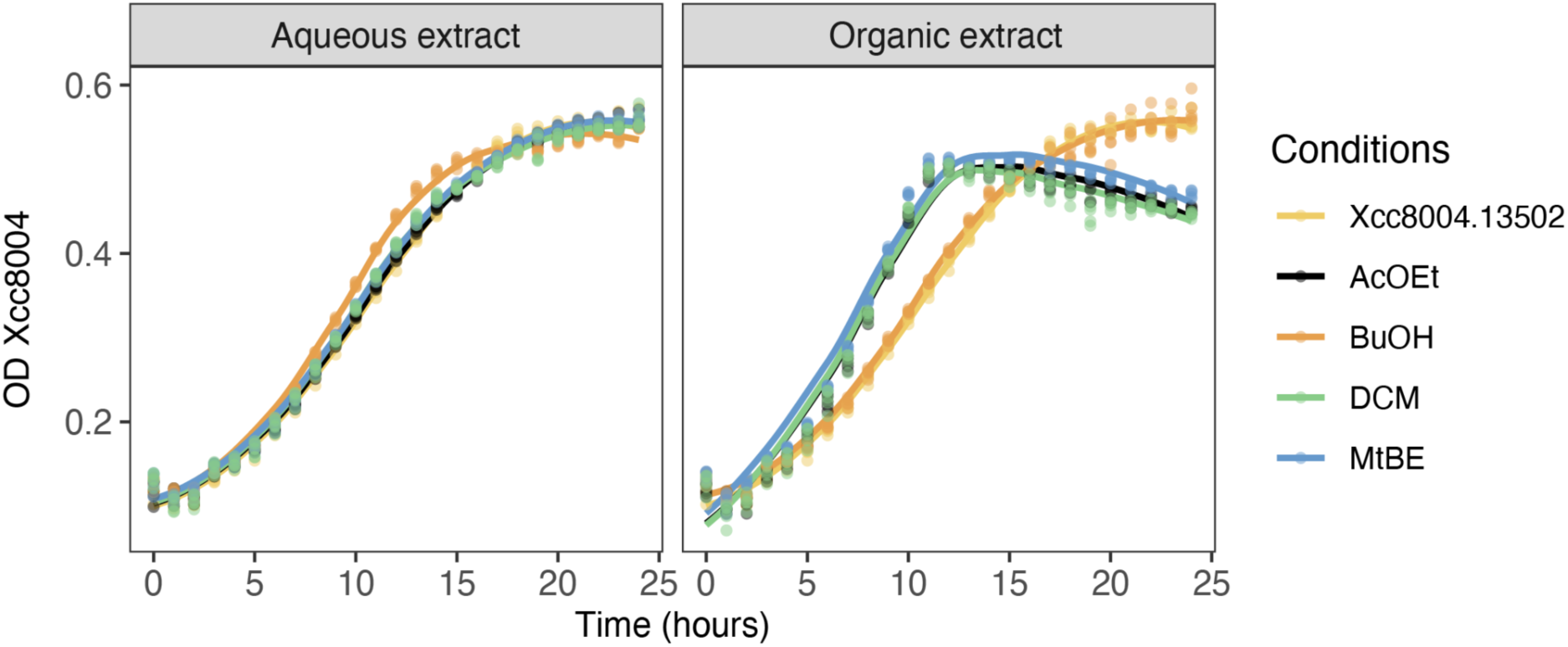
BuOH extracts inhibits Xcc8004 growth. Xcc8004 growth monitored in TSB10 medium supplemented with fractions ethyl acetate (AcOEt), butanol (BuOH), dichloromethane (DCM) and methyl *tert*-butyl ether (MtBE) obtained from CFS of Xcc8004 and CFBP13502. **A.** Growth curves of Xcc8004 in the different fractions.

**Fig. S5:**
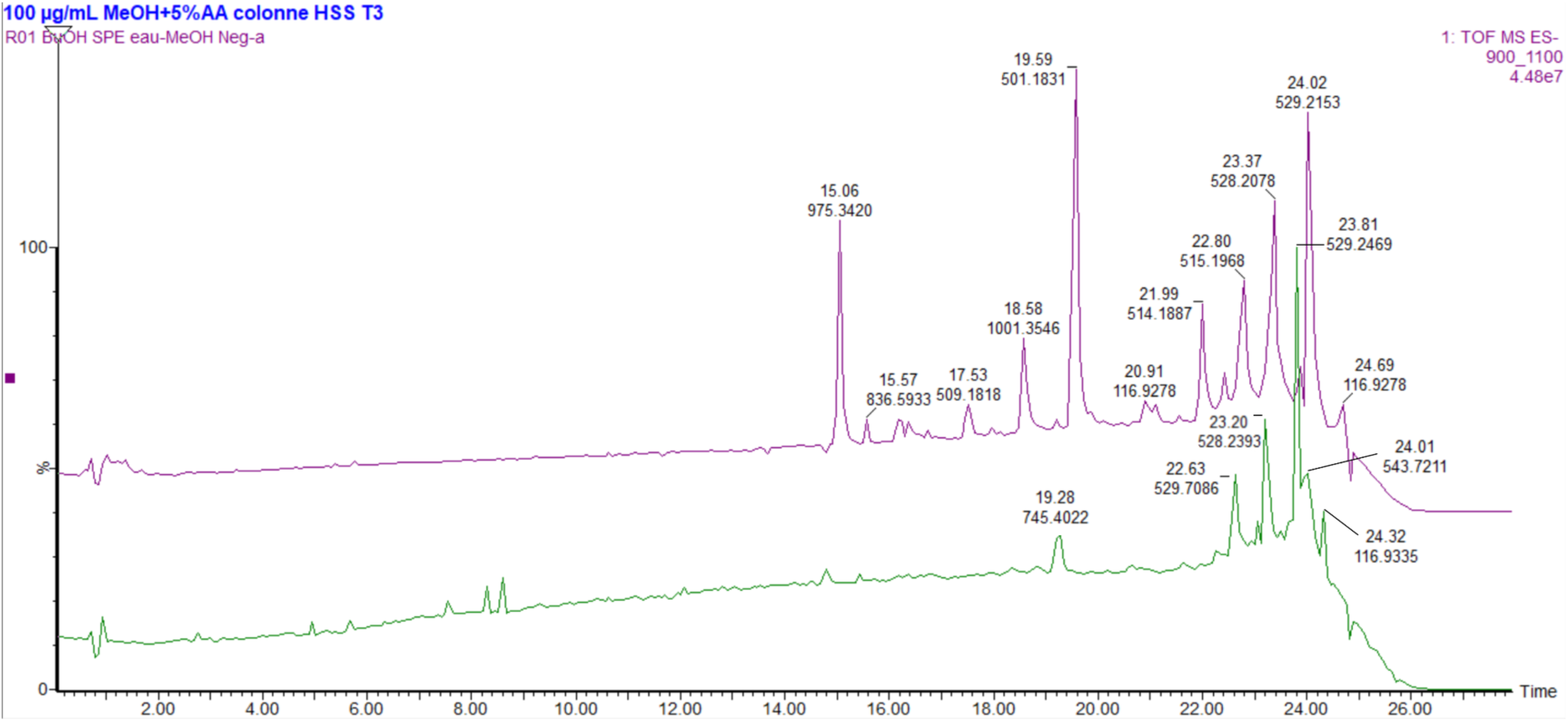
Extracted ion chromatograms of m/z between 900 and 1100, showing an increase of lactuchelins production in iron-depleted M9 medium. Green trace, bottom : CFS of coculture of CFBP13502 and Xcc8004 ; purple trace, up : single culture of CFBP13502 in iron-depleted M9 medium.

**Fig. S6:**
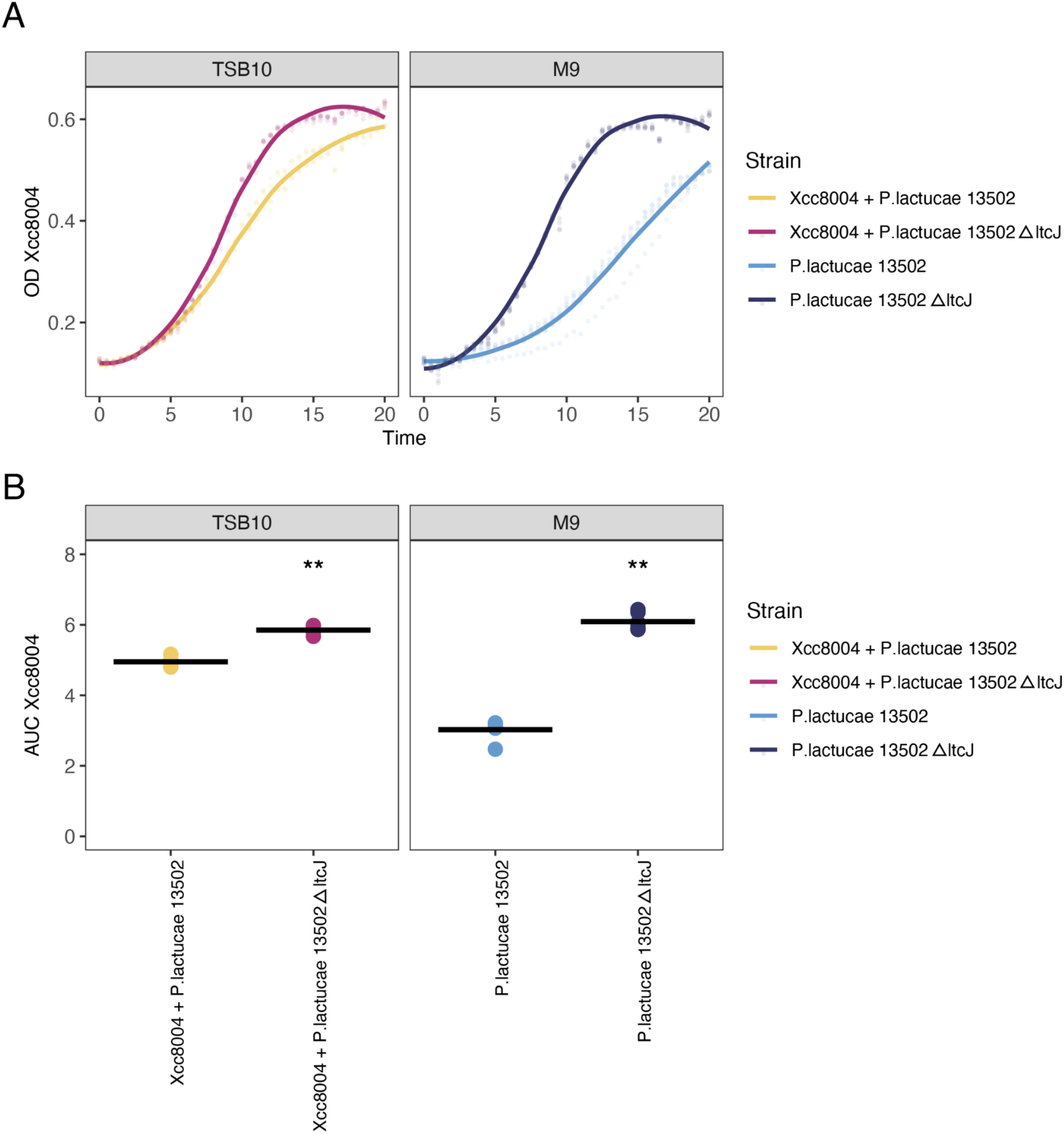
Culture of CFBP13502 in iron-depleted M9 medium improves the anti-Xcc8004 effect of the CFS. Xcc8004 growth monitored in CFSs from co-cultures of Xcc8004 and CFBP13502 (yellow), CFBP13502Δ*ltcJ* (purple) or CFBP13502Δ*ltcJ* pBBRMCS3-*ltcJ* (green). **A.** Xcc8004 growth monitored in TSB10 medium. **B.** TextTextTextTextTextText. Stars denote statistically significant differences (Wilcoxon-test, ** *P* < 0.01).

**Fig. S7:**
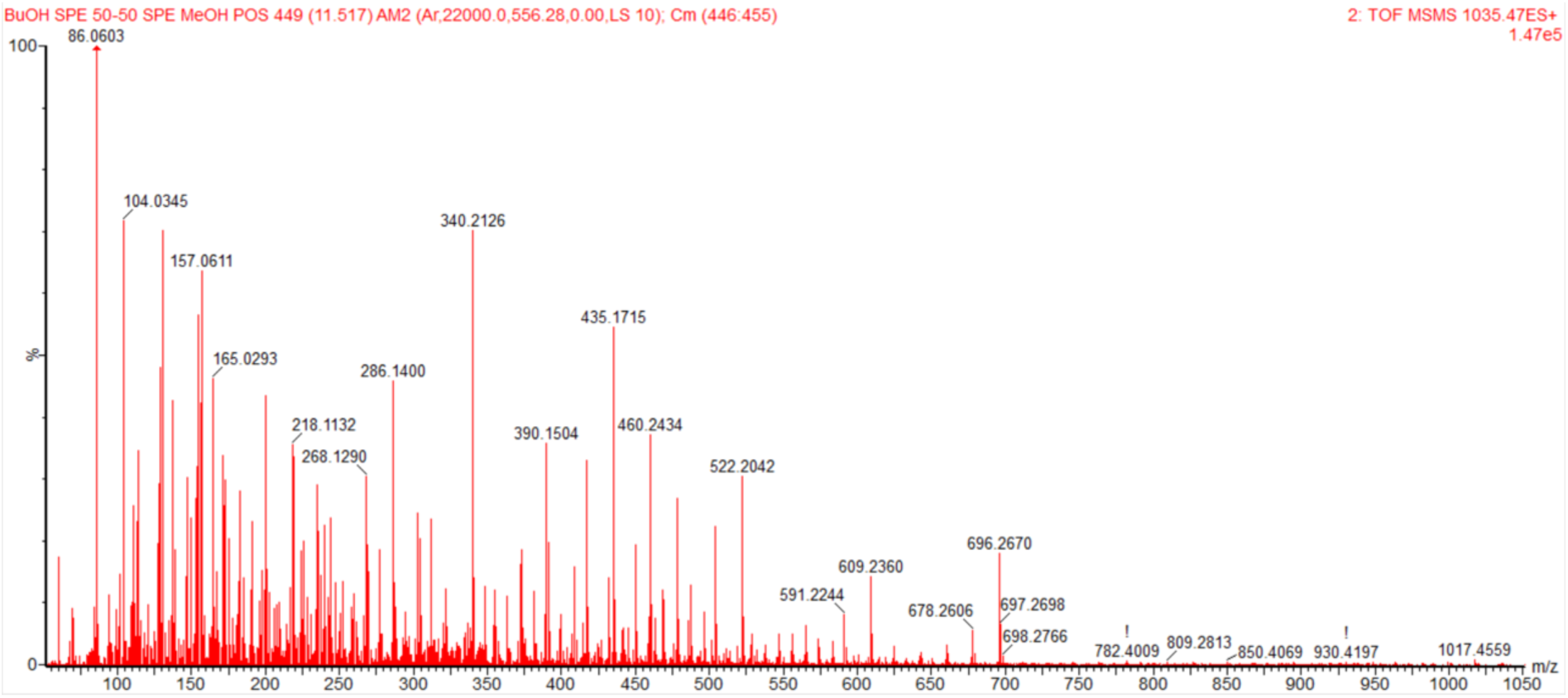
Fragmentation spectra of compound (1) in positive mode m/z 1035.4727.

## Supplementary tables

**Table S1:** Bacterial strains used in this study.

| Strain (CFBP) | Biosample | Species-Group | Host | Habitat | Year | Reference |
| --- | --- | --- | --- | --- | --- | --- |
| 8340 | SAMN41088060 | *X. campestris* pv. *campestris* 8004 | *Brassica oleracea* var. *botrytis* | unknown | 1958 | Qian et al., 2005 |
| 13502 | SAMN09044009 | *Pseudomonas lactucae* | *Raphanus sativus* Flamboyant5 | Seed | 2013 | Torres-Cortes et al., 2019 |
| 13503 | SAMN09062466 | *Stenotrophomonas rhizophila* | *Raphanus sativus* Flamboyant5 | Seed | 2013 | Torres-Cortes et al., 2019 |
| 13504 | SAMN09062476 | *Pseudomonas koreensis* subgroup | *Raphanus sativus* Flamboyant5 | Seed | 2013 | Torres-Cortes et al., 2019 |
| 13505 | SAMN09062489 | *Pantoea agglomerans* | *Raphanus sativus* Flamboyant5 | Seed | 2014 | Torres-Cortes et al., 2019 |
| 13506 | SAMN09062490 | *Pseudomonas fluorescens* subgroup | *Raphanus sativus* Flamboyant5 | Seed | 2014 | Torres-Cortes et al., 2019 |
| 13507 | SAMN09062491 | *Pseudomonas viridiflava* | *Raphanus sativus* Flamboyant5 | Seed | 2014 | Torres-Cortes et al., 2019 |
| 13508 | SAMN09062492 | *Pseudomonas koreensis* subgroup | *Raphanus sativus* Flamboyant5 | Seed | 2014 | Torres-Cortes et al., 2019 |
| 13509 | SAMN09062824 | *Pseudomonas fluorescens* subgroup | *Raphanus sativus* Flamboyant5 | Seed | 2014 | Torres-Cortes et al., 2019 |
| 13510 | SAMN09062823 | *Pseudomonas lactis* | *Raphanus sativus* Flamboyant5 | Seed | 2014 | Torres-Cortes et al., 2019 |
| 13511 | SAMN09063090 | *Erwinia persicina* | *Raphanus sativus* Flamboyant5 | Seed | 2014 | Torres-Cortes et al., 2019 |
| 13512 | SAMN09063421 | *Paenibacillus* | *Raphanus sativus* Flamboyant5 | Seed | 2014 | Torres-Cortes et al., 2019 |
| 13513 | SAMN09063410 | *Plantibacter* | *Raphanus sativus* Flamboyant5 | Seed | 2014 | Torres-Cortes et al., 2019 |
| 13514 | SAMN09063676 | *Pseudomonas fluorescens* subgroup | *Raphanus sativus* Flamboyant5 | Seed | 2014 | Torres-Cortes et al., 2019 |
| 13515 | SAMN09064882 | *Pseudomonas viridiflava* | *Raphanus sativus* Flamboyant5 | Seed | 2014 | Torres-Cortes et al., 2019 |
| 13516 | SAMN09064883 | *Pantoea agglomerans* | *Raphanus sativus* Flamboyant5 | Seed | 2014 | Torres-Cortes et al., 2019 |
| 13517 | SAMN09064884 | *Pseudomonas fluorescens* subgroup | *Raphanus sativus* Flamboyant5 | Seed | 2014 | Torres-Cortes et al., 2019 |
| 13528 | SAMN09064885 | *Pseudomonas fluorescens* subgroup | *Raphanus sativus* Flamboyant5 | Seed | 2014 | Torres-Cortes et al., 2019 |
| 13529 | SAMN09064919 | *Stenotrophomonas rhizophila* | *Raphanus sativus* Flamboyant5 | Seed | 2014 | Torres-Cortes et al., 2019 |
| 13530 | SAMN09064920 | *Enterobacter cancerogenus* | *Raphanus sativus* Flamboyant5 | Seed | 2013 | Torres-Cortes et al., 2019 |
| 13531 | SAMN16237854 | *Bacillus simplex* | *Raphanus sativus* Flamboyant5 | Seed | 2013 | Garin et al., 2024 |
| 13532 | SAMN09064921 | *Pantoea agglomerans* | *Raphanus sativus* Flamboyant5 | Seed | 2013 | Torres-Cortes et al., 2019 |
| 13533 | SAMN16237857 | *Pseudomonas orientalis* | *Raphanus sativus* Flamboyant5 | Seed | 2013 | Garin et al., 2024 |
| 13566 | SAMN16237858 | *Pantoea agglomerans* | *Raphanus sativus* Flamboyant5 | Seed | 2014 | Garin et al., 2024 |
| 13569 | SAMN16237903 | *Pantoea agglomerans* | *Raphanus sativus* Flamboyant5 | Seed | 2015 | Garin et al., 2024 |
| 13570 | SAMN16237824 | *Plantibacter* | *Raphanus sativus* Flamboyant5 | Seed | 2013 | Garin et al., 2024 |
| 13572 | SAMN16237826 | *Pseudomonas viridiflava* | *Raphanus sativus* Flamboyant5 | Seed | 2013 | Garin et al., 2024 |
| 13573 | SAMN16237813 | *Pseudomonas lactis* | *Raphanus sativus* Flamboyant5 | Seed | 2013 | Garin et al., 2024 |
| 13574 | SAMN16237900 | *Pseudomonas syringae* group | *Raphanus sativus* Flamboyant5 | Seed | 2015 | Garin et al., 2024 |
| 13575 | SAMN16237901 | *Pseudomonas coleopterorum* | *Raphanus sativus* Flamboyant5 | Seed | 2015 | Garin et al., 2024 |
| 13649 | SAMN16237913 | *Pseudomonas fluorescens* subgroup | *Raphanus sativus* Flamboyant5 | Flower | 2016 | Garin et al., 2024 |

**Table S2:**
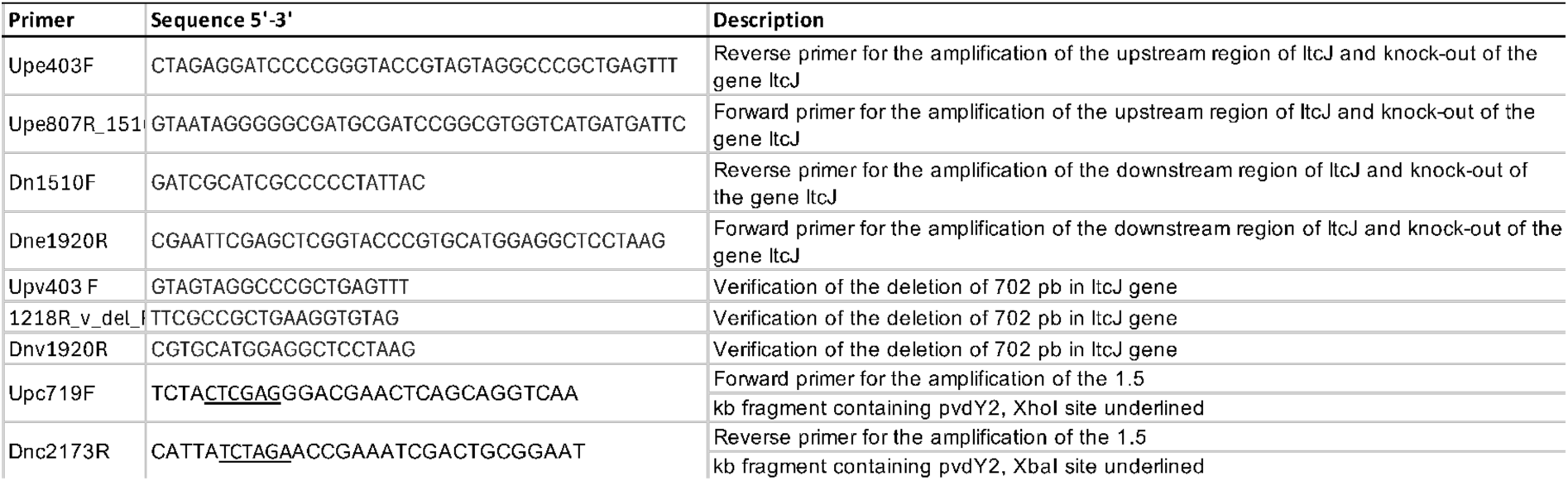
Primers used in this study.

**Table S3:**
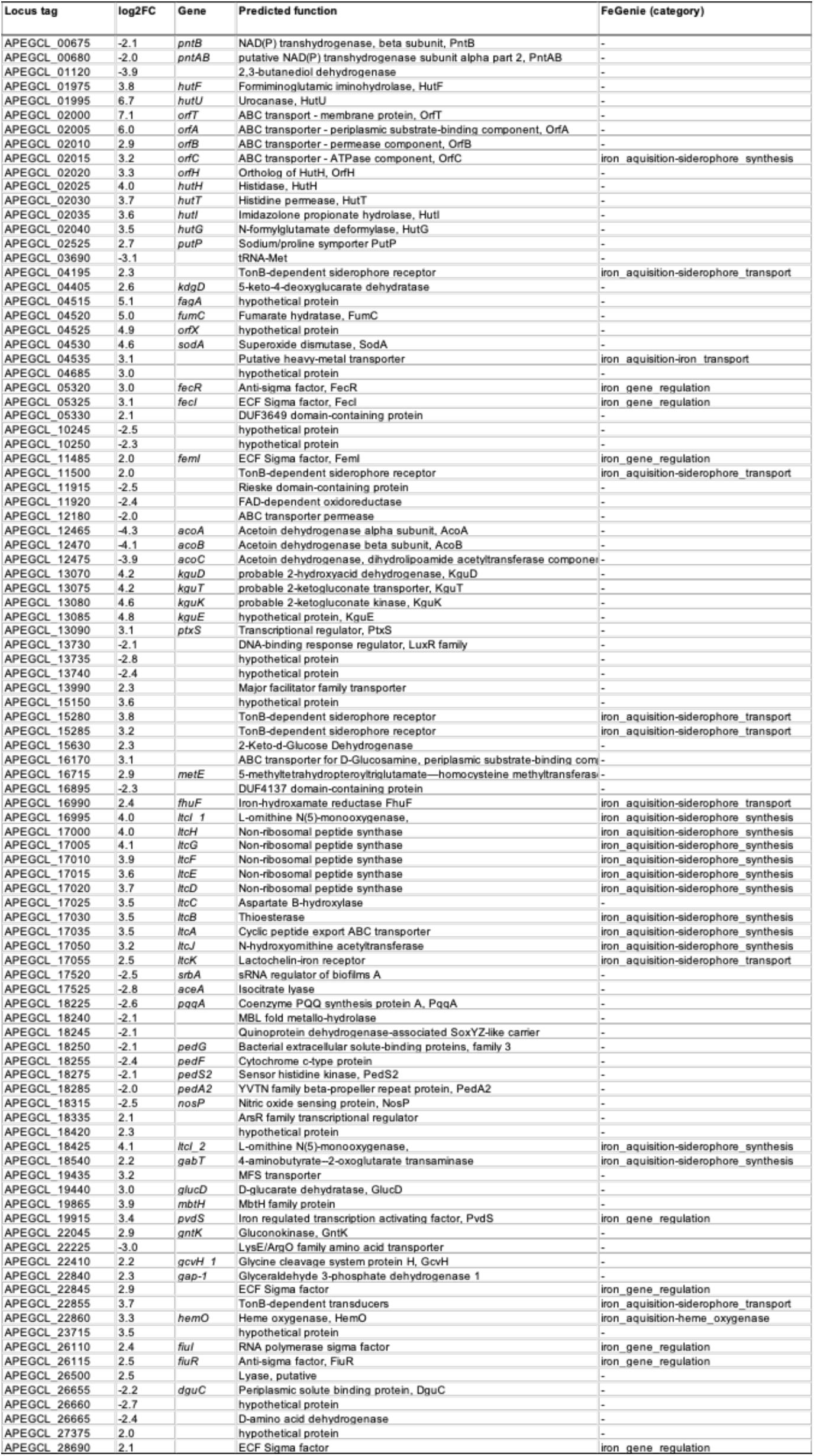
Differentially expressed genes of *P. lactucae* CFBP13502. Comparison of CFBP13502 transcriptome profiles was performed during co-cultures with (i) strains that produced a CFS with Xcc8004 inhibitory activity and (ii) strains that did not trigger CFS with inhibitory activity. Only differentially expressed genes at a *P* < 0.01 and |log_2_FC| > 2 are displayed.

**Table S4:**
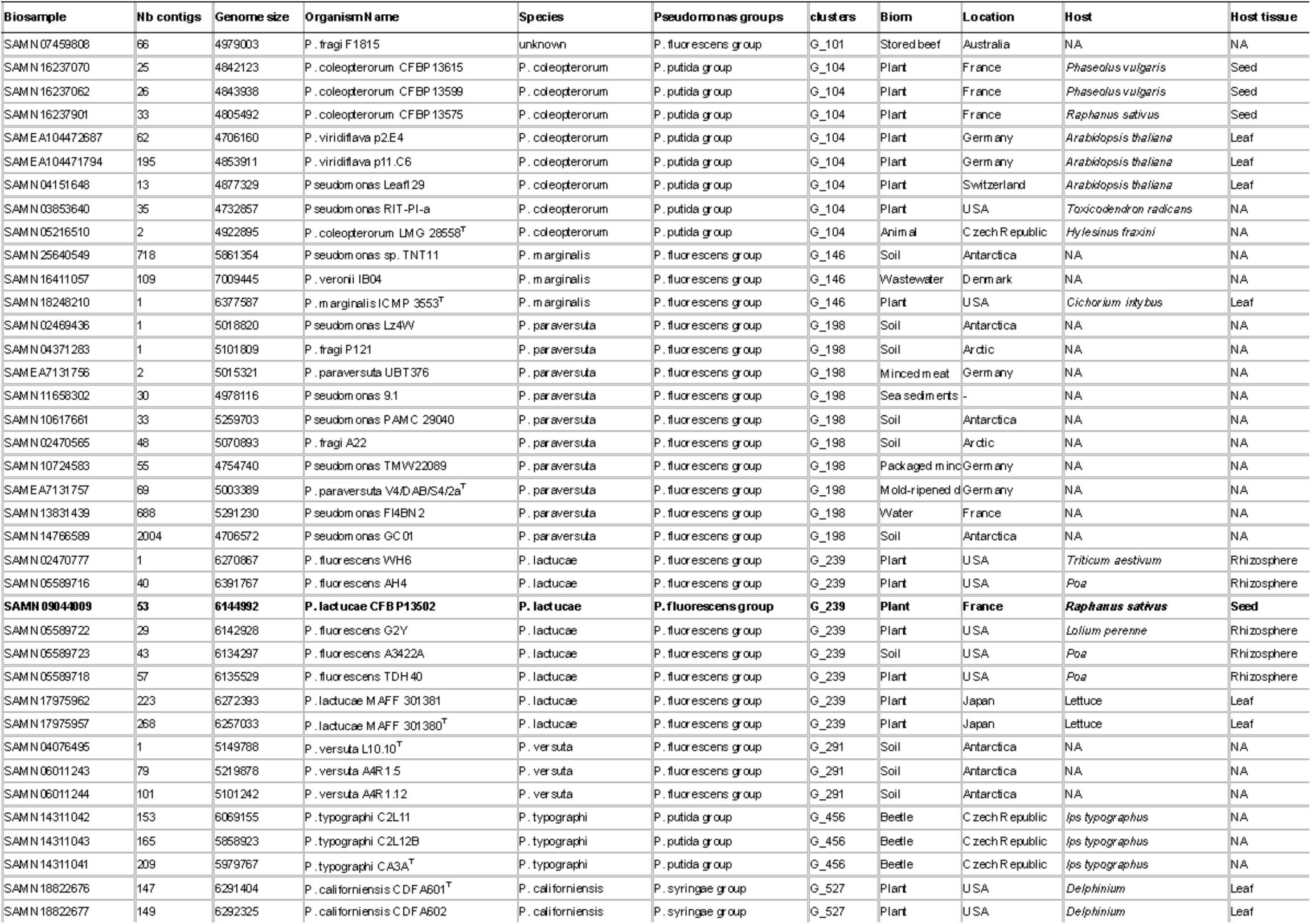
Distribution of the lactuchelin BGC within *Pseudomonas* strains.

**Table S5:**
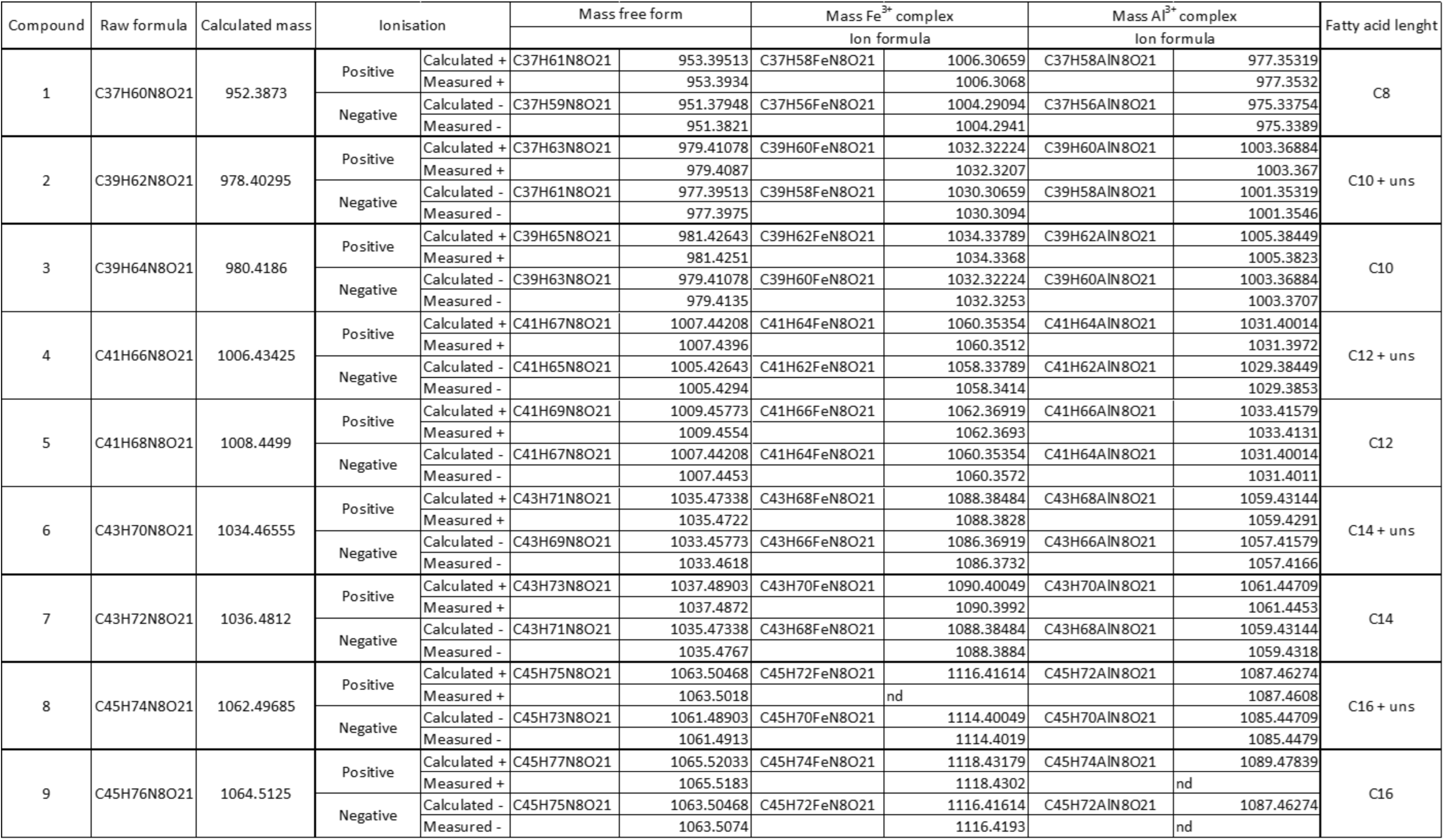
Compounds corresponding to lactuchelins detected in CFS of CFBP13502 and masses of their different forms (free, Fe3+, Al3+). nd : not detected, uns : unsaturated form

**Table S6:** ^1^H NMR (600 MHz) and ^13^C NMR (150 MHz) spectroscopic data of Ga(III)-(1) in DMSO- d_6_ (δ in ppm, J in Hz).

| No | $\delta_{\text{H}}$ mult. (J) | $\delta_{\text{C}}$ |
| --- | --- | --- |
| 1 | - | 173.6 |
| 2 | 3.82, d (2.5) | 74.2 |
| 3 | 4.43, dd (8.2; 2.0) | 56.1 |
| 4 | - | 167.9 |
| 5 | 5.37, m | 67.1 |
| 6 | (a) 2.69, dd (15.2; 11.3) | 35.3 |
|  | (b) 2.89, dd (15.2; 11.3) |  |
| 7 | - | 162.6 |
| 8 | (a) 3.39 (overlapped by residual water signal, deduced from 2D NMR) | 50.5 |
|  | (b) 4.13, m |  |
| 9 | (a) 1.65, m | 22.3 |
|  | (b) 1.83, m |  |
| 10 | (a) 1.69, m | 28.9 |
|  | (b) 2.02, m |  |
| 11 | 4.35, td (9.3; 4.4) | 52.7 |
| 12 | - | 171.4 |
| 13 | 6.88, d (8.5) | - |
| 14 | 4.19, m | 55.0 |
| 15 | (a) 3.91, d (8.2) | 62.2 |
|  | (b) 4.08, d (8.2) |  |
| 16 | - | 169.5 |
| 17 | 6.99, m | - |
| 18 | 1.12, d (6.1) | 19.8 |
| 19 | 8.05, d (8.7) | - |
| 20 | - | 169.7 |
| 21 | 4.27, m | 56.4 |
| 22 | (a) 3.65, dd (10.9; 2.6) | 60.6 |
|  | (b) 3.78, m |  |
| 23 | 7.84, d (7.3) | - |
| 24 | - | 171.8 |
| 25 | 4.08, d (8.2) | 54.8 |
| 26 | (a) 3.71, dd (10.7; 2.4) | 61.6 |
|  | (b) 3.86, dd (10.7; 2.4) |  |
| 27 | 6.99, m | - |
| 28 | - | 168.8 |
| 29 | 4.19, m | 56.14 |
| 30 | (a) 3.51, dd (10.3; 2.7) | 60.8 |
|  | (b) 3.78, m |  |
| 31 | 7.66, br. s | - |
| 32 | - | 170.1 |
| 33 | 4.71, d (8.0) | 56.5 |
| 34 | 4.61, s | 71.0 |
| 35 | - | 169.9 |
| 36 | 8.20, d (8.0) | - |
| 37 | - | 173.2 |
| 38 | 2.17, m | 35.9 |
| 39 | 1.49, m | 25.1 |
| 40 | 1.26, m | 28.3 |
| 41 | 1.26, m | 28.9 |
| 42 | 1.97, m | 26.6 |
| 43 | 5.32, m | 129.6 |
| 44 | 5.32, m | 129.6 |
| 45 | 1.97, m | 26.5 |
| 46 | 1.29, m | 29.1 |
| 47 | 1.26, m | 28.3 |
| 48 | 1.26, m | 31.2 |
| 49 | 1.26, m | 22.1 |
| 50 | 0.86, t (6.8) | 14.0 |
| 15-OH | 9.37, br. s | - |
| 34-OH | 10.99, br. s | - |

